# Imaging spatial transcriptomics reveals molecular patterns of vulnerability to pathology in a transgenic α-synucleinopathy model

**DOI:** 10.1101/2024.07.31.606032

**Authors:** Liam Horan-Portelance, Michiyo Iba, Dominic J. Acri, J. Raphael Gibbs, Mark R. Cookson

## Abstract

In Parkinson’s disease and dementia with Lewy bodies, aggregated and phosphorylated α-synuclein pathology appears in select neurons throughout cortical and subcortical regions, but little is currently known about why certain populations are selectively vulnerable. Here, using imaging spatial transcriptomics (IST) coupled with downstream immunofluorescence for α-synuclein phosphorylated at Ser129 (pSyn) in the same tissue sections, we identified neuronal subtypes in the cortex and hippocampus of transgenic human α-synuclein-overexpressing mice that preferentially developed pSyn pathology. Additionally, we investigated the transcriptional underpinnings of this vulnerability, pointing to expression of *Plk2*, which phosphorylates α-synuclein at Ser129, and human *SNCA* (*hSNCA*), as key to pSyn pathology development. Finally, we performed differential expression analysis, revealing gene expression changes broadly downstream of *hSNCA* overexpression, as well as pSyn-dependent alterations in mitochondrial and endolysosomal genes. Overall, this study yields new insights into the formation of α-synuclein pathology and its downstream effects in a synucleinopathy mouse model.

## Introduction

One of the shared neuropathological hallmarks of Parkinson’s disease (PD) and dementia with Lewy bodies (DLB) is Lewy pathology, which consists of misfolded, aggregated, and phosphorylated forms of the protein α-synuclein localized to the cell bodies (Lewy bodies, LBs), neurites (Lewy neurites) and synapses of affected neurons^1–4^. The presence of Lewy pathology may be pathogenic, as evidence points to transcriptional and functional deficits of afflicted neurons^5, 6^. Lewy pathology may also precede degeneration of the cells it affects, as evidenced by time-course experiments tracking the appearance of pathology and subsequent degeneration of neurons in the same brain regions in mice^7, 8^. However, multiple reports of familial variants of PD which occur in the absence of Lewy pathology^9–11^ and individuals with abundant Lewy pathology but without neurocognitive deficits^12, 13^ leave open the question as to whether Lewy pathology is a pathogenic driver of disease or part of a response to other disease mechanisms^14^.

Across neurodegenerative diseases, neurons display selective vulnerability to disease-associated pathologies, degeneration, or both; this vulnerability is apparent at the brain region and neuronal subtype level^15^. In PD, Lewy pathology first emerges in deep subcortical regions, progressing to the midbrain and finally higher-order neocortical regions^16^. The presence of midbrain Lewy pathology is accompanied by the loss of dopaminergic neurons (DANs) in the substantia nigra pars compacta (SNc), which drives the disease’s classical motor symptoms^17^. In DLB, the progression of pathology differs, with faster progression in limbic and cortical regions^18^. In both diseases, neurons show patterns of selective vulnerability to Lewy pathology; in PD, SNc DANs develop abundant pathology while adjacent DANs of the ventral tegmental area (VTA) are relatively resistant^19^, and in DLB, neuritic Lewy pathology develops in the CA2 hippocampal subfield, while the dentate gyrus is spared^8, 20^. In the neocortex, excitatory neurons (ExNs) develop LBs, while inhibitory neurons (InNs) do not^5, 21, 22^, and intratelencephalic neurons of layer 5 (L5 IT), a subtype of pyramidal ExNs, may be more vulnerable to Lewy pathology relative to other ExN subtypes^5^.

Despite an understanding of which regions and cell types are vulnerable to pathology and degeneration, the underlying molecular mechanisms of vulnerability remain to be fully described. Some evidence implicates endogenous α-synuclein expression as a driver of vulnerability to Lewy pathology^8, 23^. For example, ExNs, which develop Lewy pathology, express high levels of endogenous α-synuclein, and InNs and non-neuronal cells, which are resistant, express much lower levels^21, 22, 24^. However, several observations indicate that endogenous α-synuclein expression does not fully determine cellular vulnerability to pathology and degeneration in synucleinopathies. For example, SNc and VTA DANs both express very high endogenous α-synuclein, but have very different vulnerability, to both pathology and degeneration^19, 24^. Additionally, in multiple system atrophy (MSA), α-synuclein accumulates in oligodendrocytes, despite these cells exhibiting little to no basal α-synuclein expression^25, 26^. Other work has suggested that factors such as functional connectivity, myelination state, degree of axonal branching, and calcium buffering influence neurons’ vulnerability, but these links have not been established as causal^19, 21, 27^. Recent work using single-cell/nucleus RNA sequencing (sc/snRNA-seq) in the context of tauopathy suggests that there are innate transcriptional differences between vulnerable and resilient neurons which may help explain their vulnerability to tau pathology^28, 29^. For example, InNs, which largely do not develop tau pathology, express higher levels of genes in pathways involved in autophagic processing of tau, which could contribute to their resilience^29^. Similar transcriptional differences may exist in synucleinopathies, which could help explain selective vulnerability to α-synuclein pathologies where α-synuclein expression alone fails to do so, which we sought to address for the first time in the present study.

Recent advances in spatial transcriptomic technologies have enabled imaging-based detection of hundreds of genes simultaneously at subcellular resolution. Since these technologies are non-destructive, they can be coupled with downstream analyses, including immunofluorescence (IF) for protein co-detection in the same sections^30^. In the current study, we used Xenium, a subcellular-resolution imaging spatial transcriptomic (IST) platform, with a well-characterized transgenic synucleinopathy mouse model which overexpresses wild-type human α-synuclein (*hSNCA*) under the murine neuronal Thy1 promoter (Line 61, α-syn-tg). By performing post-Xenium IF with an antibody against α-synuclein phosphorylated at Serine 129 (pSyn), we identified cortical and hippocampal cell types that were vulnerable and resilient towards developing this pathology. Next, by creating a custom panel of probes against genes known to be involved in α-synuclein processing, aggregation, and toxicity, we interrogated the transcriptional underpinnings of the selective neuronal vulnerability to α-synuclein pathology in this model. Finally, by independently comparing transcriptomes of healthy non-transgenic (non-tg) neurons to pSyn^+^ and pSyn^-^ α-syn-tg neurons, we could distinguish conserved transcriptional changes associated generally with *hSNCA* overexpression, such as molecular chaperones and autophagy-related genes, versus those which were pSyn pathology-dependent, including genes related to mitochondria and the endolysosomal system. This study uses novel IST technology to yield new biological insights into the neuronal vulnerability to and downstream effects of α-synuclein pathology in this α-syn-tg mouse model.

## Methods

### Mouse line

Line 61 (α-syn-tg) mice were originally generated by inserting cDNA for the coding region of human *SNCA* under the control of the mouse *Thy1* promoter^31^ and is available from JAX (strain #038796). Eight 7-month-old mice were used, 4 non-tg and 4 α-syn-tg, with 2 males and 2 females of each genotype. See Supplementary Table 1 for specific information on each animal. All animal experiments were carried out in accordance with protocol 463-LNG-2024, approved by the Institutional Animal Care and Use Committee (IACUC) of the National Institute on Aging (NIA).

### Mouse brain sample preparation

Mice were harvested via trans-cardiac perfusion with phosphate-buffered saline (PBS) after injection of pentobarbital. Brains were removed from the skull, and the right hemispheres were fixed in 10% neutral buffered formalin (NBF) overnight at 4°C, cut into 4 mm thickness coronally, dehydrated, then embedded in paraffin. Four hemispheres of non-tg or α-syn-tg were embedded together in one block, then sections were cut at a thickness of 6 μm on a microtome and mounted onto Xenium slides or normal glass slides.

### Xenium *in situ* transcriptomics

Xenium sample preparation protocol was carried out according to manufacturer’s instructions, which are available as 10x Genomics Demonstrated Protocols, including Tissue Preparation (CG000578), Deparaffinization and Decrosslinking (CG000580), and Xenium In Situ Gene Expression (CG000582), and in Janesick et al., 2023^32^. Briefly, sections were deparaffinized and rehydrated using a xylene and ethanol series, followed by decrosslinking to facilitate mRNA availability. Probes were hybridized overnight at 50°C (full list of base and custom panel probes can be found in Supplementary Table 2). Following washing to remove unbound probes, probes were ligated for 2 hours at 37°C, followed by enzymatic rolling circle amplification for 2 hours at 30°C. Finally, autofluorescence quenching was performed followed by nuclei staining, and slides were loaded onto the Xenium Analyzer instrument (software v1.7.6.0). On the instrument, samples underwent successive rounds of fluorescent probe hybridization, imaging, and probe removal.

### Post-Xenium immunofluorescence (IF)

Following completion of the Xenium run, slides were stored in PBST at 4°C in the dark until use. We performed antigen retrieval with citrate-based buffer (Vector, H-3300) and microwave heat. Slides were blocked in 4% fetal bovine serum, followed by incubation with primary antibodies (pSer129 α-synuclein, clone EP1536Y, abcam, ab51253, 1:2000; NeuN, Millipore, MAB377, 1:500) overnight in blocking buffer at 4°C. The following day, slides were washed with PBS and incubated again with corresponding secondary antibodies (goat anti-rabbit-Texas Red, TI-1000; horse anti-mouse-Fluorescin, FI-2000, both Vector) overnight at 4°C. On the third day, slides were washed, incubated with DAPI (Invitrogen, H3569) for 20 minutes, and autofluorescence was quenched with TrueBlack Lipofuscin Quencher (Biotium, 23007). Finally, slides were coverslipped with ProLong Gold antifade reagent (Thermo Fisher, P36930). Whole-tissue scans were acquired on a LSM780 confocal microscope (Zeiss) in the red and blue channels, for pSyn and DAPI, respectively. Tiled images were acquired at 20x magnification with 10% overlap at a resolution of 1024 x 1024 pixels, with the same laser and gain settings across sections. For representative images (Supplementary Fig. 1), images were acquired at 10x or 40x in a single field of view in the red and green channels, for pSyn and NeuN, respectively. For both whole-slide and representative images, colors were changed using the lookup tables function in Fiji (NIH), and for representative images, brightness and contrast were altered, the same across each image.

### Xenium data processing

Xenium data was re-segmented using Xenium Ranger (v1.7.1.1, 10x Genomics), changing from a 15 to a 5 μm nuclear expansion method. Nuclei were first identified and segmented through the DAPI stain, after which, cell boundaries were expanded 5 μm or until they encounter another cell’s boundaries. Data was then loaded into Seurat^33^ (v5.0+) and processed using standard scRNA-seq workflows. Briefly, counts were normalized against library size and log-transformed using NormalizeData(); data was scaled with ScaleData(), regressing out number of reads per cell; PCA was computed using RunPCA(); finally, datasets were integrated using Harmony^34^. Cell barcodes from the cortex or hippocampus of each sample were determined by manually drawing regions of interest within Xenium Explorer (v2.0+, 10x Genomics), and the Seurat object was subset accordingly to only contain either cortical or hippocampal cells. After subsetting, data was re-scaled for both cortex and hippocampus independently, followed by the standard Seurat single-cell workflow (RunPCA(), RunUMAP(), FindNeighbors(), and FindClusters()). Due to the relative differences in cell type diversity between the two regions, clustering resolution was set to 0.8 for the cortex and 0.3 for the hippocampus to avoid over-clustering. Cluster identification was performed manually using marker genes determined using FindAllMarkers() in Seurat and cross-referencing against canonical cell type markers.

### Xenium/pSyn overlap analysis

After IF image acquisition, .czi images were converted to pyramidal OME TIFFs using QuPath^35^. Using Xenium Explorer, IF images were aligned to and overlaid with Xenium data. Affine transformation matrices of geometric translations and transformations used to align IF images to Xenium data in Xenium Explorer were extracted. IF images were loaded back into QuPath, and the same affine transformations were applied, yielding a transformed image on the same coordinates as those of the Xenium data. Raw and transformed images, as well as affine transformation matrices generated by Xenium Explorer, are available for download on Zenodo. Using QuPath, IF images were thresholded in the pSyn channel using the same thresholds across all four α-syn-tg brains, and the coordinates of the centroids of the thresholded pSyn inclusions were obtained. In R, we used the sf package^36, 37^ to overlay the centroids of the pSyn inclusions with the polygons of the cells from the Xenium data; if the centroid of a pSyn inclusion fell within the bounds of a Xenium cell’s polygon, that cell was considered pSyn^+^. pSyn positivity information was added to Seurat metadata of the Xenium object and used for downstream analysis.

### Comparison of Xenium and single-cell RNA sequencing data

We compared our Xenium data to the Allen Brain Cell (ABC) atlas isocortex data^38^. ABC atlas isocortex data was downloaded (10x v2 chemistry) and downsampled to 240,000 total cells. Raw counts were normalized to library size in Seurat using LogNormalize() with a scale factor of 10^6^. We then subset the ABC atlas object to only include genes which were present in our base or custom Xenium panels. To facilitate direct comparison of expression data between Xenium and the ABC atlas data, gene expression was averaged (using AverageExpression() in Seurat) and scaled across the selected cell types. For correlating expression of individual genes in the base or custom panels, a linear regression was run for each gene, correlating the scaled expression of that gene in the defined cell types in the ABC atlas against its expression in the Xenium data in those same cell types.

For comparing cell type gene expression to validate cell type assignments, we first generated “expression profiles” for each cell type in the ABC atlas and Xenium datasets by scaling the average expression of each gene in the base Xenium panel across the same cell types in both datasets independently. We then computed a Pearson coefficient between the expression profiles of ABC atlas and Xenium cell types, obtaining the similarity score between all the cell types from both datasets. Note that for both these analyses (i.e., comparing expression of individual genes and expression profiles of whole cell types), only non-tg cells were used from the Xenium experiments, as to avoid confounds of gene expression changes in α-syn-tg cells.

### Pseudo-bulk differential expression analysis

DESeq2^39^ was used for pseudo-bulk differential expression analysis. Pseudo-bulk counts for the cortex were first aggregated across cell type, sample, and pSyn status using AggregateExpression in Seurat, followed by standard DESeq workflows. DESeq2 automatically corrects for biological replicates, so our design formulas did not include sample as a covariate. Only genes from the custom add-on panel were considered, and genes were considered significantly differentially expressed if the Benjamini-Hochberg (B-H) adjusted p-value (FDR) was < 0.05. For analysis across multiple cell types, we only considered genes which were significant DEGs in the same direction (i.e., upregulated or downregulated) in 2 or more cell types. Additionally, since hippocampal differential expression is known to be affected by position on the rostro-caudal axis^40, 41^, and given the very low number of pSyn^+^ inhibitory neurons in this model, we did not include these cell types in our differential expression analysis, restricting it to solely excitatory neurons in the cortex.

### Generalized linear modeling (GLM) of gene expression against *hSNCA*

Monocle3^42–45^ was used to perform GLM analysis, in which we regressed the expression of individual genes of interest (gene expression, GEx) against expression of *hSNCA* in single cells, aiming to determine, for a given cell type, if the expression of a gene of interest was correlated with *hSNCA* expression. To facilitate this, *hSNCA* expression was added to the cell-level metadata of the Seurat object. α-Syn-tg cells from each ExN subtype of interest were subset from the Seurat object, and a CellDataSet (cds) object was created using Moncle for each subtype. Each cds was preprocessed using Monocle’s preprocess_cds() function with 30 dimensions, and the GLM was run using the fit_models() function with the model “GEx ∼*hSNCA* expression”. Only genes which were found to be differentially expressed using the pseudo-bulk method were considered for the GLM analysis. GLM results were considered significant if the B-H adjusted p-value (i.e., q-value) was < 0.05.

### Immunofluorescence for proteinase K-resistant α-synuclein

To visualize proteinase K (PK)-resistant α-synuclein, we sacrificed non-tg and α-syn-tg as described above and post-fixed with 4% paraformaldehyde. Paraffin sections were prepared the same way as for Xenium, and the same immunofluorescence protocol was followed, except that after antigen retrieval, slides were incubated with 1 ug/mL PK (Viagen, 501-PK) for 1 minute at room temperature. We stained with an antibody against total α-synuclein (BD Biosciences, 610787), which was detected using donkey anti-mouse AlexaFluor 647 (Invitrogen, A-31571). Images were acquired on a Nikon CSU-W1 SoRa confocal microscope using the same settings across all images. Images were processed using Fiji, with the same brightness/contrast alterations applied across all images.

### Statistics

For comparing pSyn pathology frequency across cell types, we performed a one-way ANOVA with Tukey’s multiple comparisons test using GraphPad Prism (v10+). To correlate average gene expression in a cell type against pSyn pathology rates in that same cell type, we used a linear regression using the lm() function in R with the model “GEx ∼% cells pSyn^+^”. Raw p-values were corrected for multiple comparisons using the Benjamini-Hochberg method in GraphPad Prism. As stated above, correlating gene expression against hSNCA expression was performed using Monocle3 in R, using a GLM. Pseudobulk differential gene expression was performed using DESeq2 as described above, which assumes a negative binomial distribution of counts data, and uses a GLM with the Wald test for significance for generating DE results.

## Results

In the present study, we used a well-characterized mouse model of α-synucleinopathy (Line 61, α-syn-tg) which overexpresses wild-type human α-synuclein (*hSNCA*) under the murine Thy1 promoter, and non-transgenic littermate controls (non-tg). α-Syn-tg mice develop abundant α-synuclein pathology throughout cortical and subcortical brain regions and display PD-relevant motor and non-motor behavioral deficits^31, 46, 47^. Neurons in this model develop intracellular α-synuclein inclusions, which we detected in this experiment using an antibody against α-synuclein phosphorylated at serine 129 (Supplementary Fig. 1a). Notably, these mice have distinct phospho-α-synuclein pathology^48^, including nuclear staining (Supplementary Fig. 1b), and in severely afflicted neurons, cytoplasmic and axonal pathology (Supplementary Fig. 1c). Line 61 α-syn-tg mice also display accumulation of proteinase-K (PK)-resistant α-synuclein in the cell bodies of some neurons (Supplementary Fig. 2a-c).

Coronal formalin-fixed, paraffin embedded (FFPE) brain sections from non-tg and α-syn-tg mice (n = 4 each) were processed and run on Xenium followed by immunofluorescence for pSyn in the same sections, allowing us to identify cells which were pSyn^+^ or pSyn^-^. For this experiment, we analyzed pSyn pathology distribution in the neocortex and hippocampus of the mice, given the pathology burden observed in these brain regions in late-stage PD and in DLB^16^, and the relevance of these regions to cognitive symptoms in disease^13^.

### Imaging spatial transcriptomics identifies within-layer cortical cell types at single-cell resolution

Across all samples, Xenium identified 136,276 cortical cells that were segmented using a 5-μm nuclear expansion method (Fig. 1a, b). We filtered out any cells which did not contain any detected genes, and validated quality control metrics of the remaining cells (Supplementary Fig. 3a-d). We found that non-tg and α-syn-tg samples contained similar numbers of detected cells, and that cells contained similar numbers of unique genes detected per cell and low negative probe rates across genotypes (Supplementary Fig. 3a, c, d). Number of detected transcripts per cell was slightly higher in α-syn-tg cells, possibly due to detection of the transgene (Supplementary Fig. 3b).

**Fig. 1.**
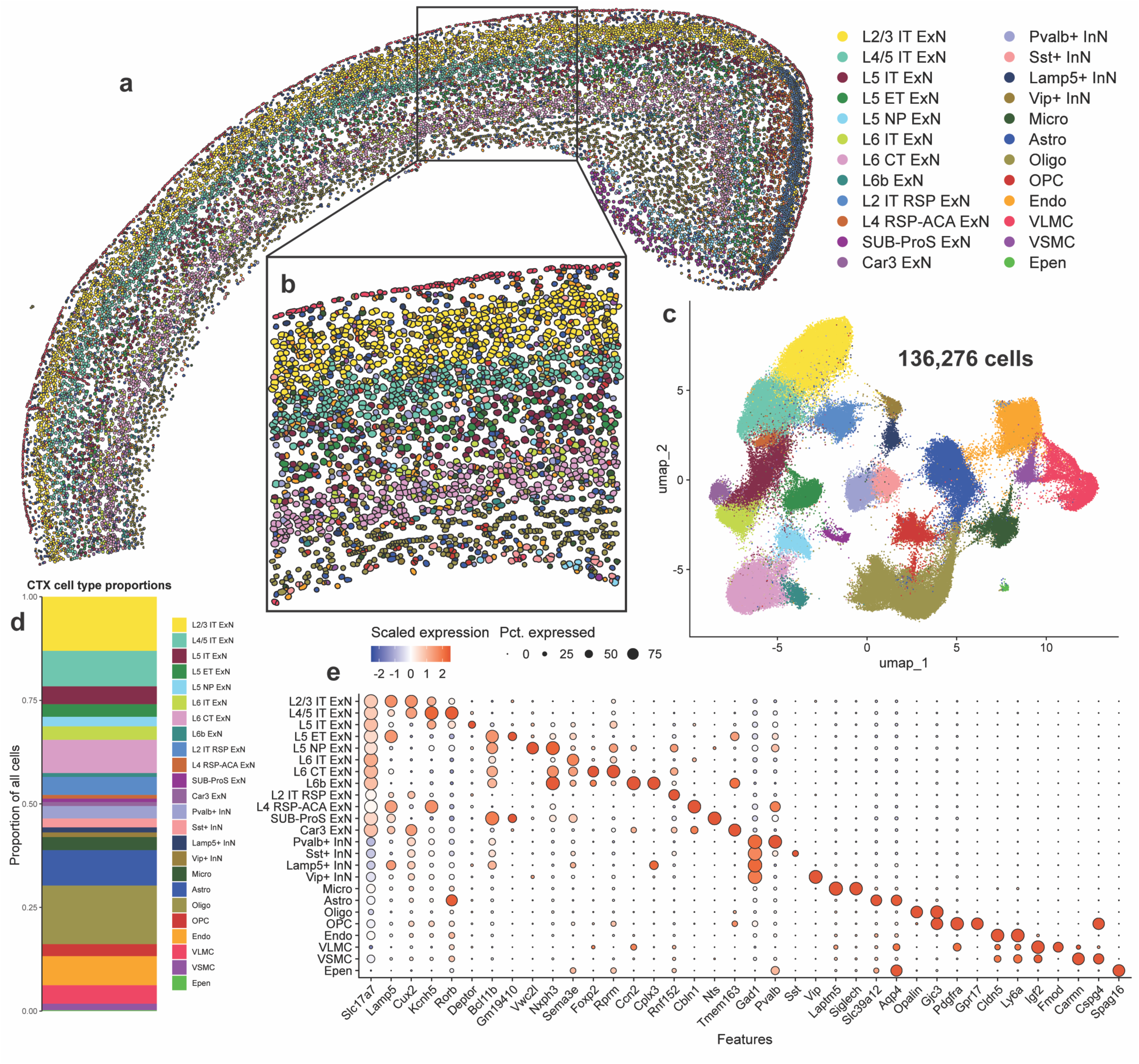
Imaging spatial transcriptomics identifies within-layer cortical cell types at single-cell resolution. **a** Cortical cells from one representative non-tg sample shown in physical space. Dots represent individual cells, colored by cell type. **b** Enlarged image showing cortical layering and cell segmentation via 5-μm nuclear expansion. Dots represent individual cells, colored by cell type. **c** All cortical cells from 8 samples (4 non-tg and 4 in α-syn-tg) shown in UMAP space (136,273 total cells). Dots represent individual cells, colored by cell type (same colors as in **a-b**). **d** Proportions of all identified cortical cell types across all 8 samples. **e** Dotplot showing scaled expression of canonical marker genes for each cortical cell type, sized by percentage of cells expressing a given marker.

We identified 24 cell clusters within the cortex, including excitatory neurons (ExNs), inhibitory neurons (InNs), glial cells, and vascular cells (Fig. 1c, e). We resolved 12 subtypes of ExNs, identified broadly by their expression of *Slc17a7*, the gene encoding vesicular glutamate transporter 1 (Vglut1); these subtypes included general ExNs, which were present in all brain sections, regardless of position on the rostro-caudal axis, and region-specific ExNs, the abundance of which varied based on section depth (Fig. 1a, b, d, e, Supplementary Table 3). These region-specific subtypes included L2 IT RSP, L4 RSP-ACA, SUB-ProS, and Car3 ExNs (Fig. 1a, c, d). For downstream analysis involving ExNs, we only considered general ExN subtypes, because region-specific ExNs were not present in representative numbers in each sample (Supplementary Table 3).

We identified 8 transcriptionally distinct subtypes of general ExNs, assigned using expression of canonical subtype-specific genes^49^ (Fig. 1e). From the outer cortical layers, we identified L2/3 intratelencephalic (IT) ExNs, which expressed high levels of *Lamp5* and *Cux2*, and L4/5 IT ExNs, which specifically expressed *Kcnh5* and *Rorb* (Fig. 1e). Within layer 5, we identified 3 subtypes which clustered independently based on their projection type; these included L5 IT, which expressed *Deptor*, L5 extratelencephalic (ET), which expressed *Bcl11b* and *Gm19410*, and L5 near-projecting (NP), which were marked by *Vwc2l* (Fig. 1e). We also identified 3 subtypes of layer 6 neurons, including L6 IT, which expressed *Sema3e*, L6 cortico-thalamic (CT), which expressed *Foxp2* and *Rprm*, and L6b, which expressed *Ccn2* and *Cplx3* (Fig. 1e). We also identified 4 subtypes of InNs, broadly defined by their expression of *Gad1*, encoding glutamate decarboxylase 1, which were split by their expression of canonical interneuron markers, namely parvalbumin (*Pvalb*), somatostatin (*Sst*), *Lamp5*, and vasoactive intestinal peptide (*Vip*) (Fig. 1e). Finally, we resolved 8 types of non-neuronal cells, including 4 glial subtypes (microglia – *Laptm5*, *Siglech*; astrocytes – *Aqp4*, *Slc39a12*; oligodendrocytes – *Opalin*; oligodendrocyte precursors – *Pdgfra*) and 4 vascular cell types (endothelial cells – *Cldn5*, *Ly6a*; vascular leptomeningeal cells (VLMC) – *Igf2*, *Fmod*; vascular smooth muscle cells (VSMC) – *Carmn*, *Cspg4*; ependymal cells – *Spag16*) (Fig. 1e).

Since IST uses fluorescence microscopy rather than sequencing to measure gene expression, we compared expression of individual genes in our Xenium-detected cells to expression of those same genes in cells from a single-cell RNA sequencing (scRNA-seq) dataset, specifically mouse isocortex from the Allen Brain Cell (ABC) atlas^38^ (Supplementary Fig. 4a, b). Given the technical differences between modalities, it is not possible to directly compare raw or normalized gene count values, so we averaged and scaled the expression of each gene across the cell types shared between the two datasets. We then performed linear regression between scaled Xenium and scRNA-seq gene expression data, separately analyzing genes in the base and custom Xenium panels (Supplementary Fig. 4a, b). We found that scaled expression of genes from both the base and custom Xenium panels correlated well (r^2^ = 0.685 and 0.418, p < 0.0001 for base and custom panels, respectively) between Xenium and scRNA-seq (Supplementary Fig. 4a, b). Most of the outlier genes in this analysis were genes expressed in non-neuronal cells, likely due to the inability of our 5-μm nuclear expansion segmentation method to capture the greater diversity of cell body shapes and sizes of these cell types compared to neurons, leading to more frequent mis-assignment of reads from neighboring cells to these non-neuronal cells than vice versa. However, our ability to correlate expression of individual genes by two orthogonal approaches indicates that neuronal transcriptional responses can be adequately monitored by IST.

Additionally, since imaging spatial transcriptomics uses a defined set of probes for cell type identification, we sought to validate our cell type assignments in the cortex, again using the ABC atlas as a reference (Supplementary Fig. 4c). Using the scaled expression of the Xenium base panel genes in the various cell types shown in Supplementary Fig. 4a, we ran a Pearson correlation between the full expression profiles of each cell type in both Xenium and the ABC atlas (Supplementary Fig. 4c). Pearson R values ranged from 0.812 to 0.942 for matched neuronal subtype pairs in Xenium and scRNA-seq data, indicating very high agreement of gene expression profiles generated by the two technologies (Supplementary Fig. 4c).

We also sought to use the ground truth of spatial localization to further validate our cell type assignments, particularly ExNs restricted to specific cortical layers. We used two representative fields of view through the whole cerebral cortex and corpus collosum, from one non-tg and one α-syn-tg animal, and calculated the localization of the various cell types as a function of depth through the cortex (Supplementary Figs. 5a-b, 6a-b). We found that spatial distribution of many cell types was strikingly restricted, corresponding with their known location in the cortex. All cortical layer-specific ExN subtypes were enriched in their expected spatial niches (Supplementary Figs. 5a-b, 6a-b). Additionally, Lamp5+ and Vip+ InNs showed the highest abundance in layer 1 (L1) of the cortex, which is known to be sparsely populated with interneurons^50^ (Supplementary Figs. 5a-b, 6a-b). Finally, some non-neuronal cells showed spatial specificity, namely VLMCs, which were found at the outer edges of the cortex, and oligodendrocytes, which were primarily found in the corpus collosum. In contrast, but as expected, microglia, astrocytes, and endothelial cells were evenly distributed throughout the cortex (Supplementary Figs. 5a-b, 6a-b).

Taken together, these data demonstrate that Xenium imaging-based spatial transcriptomics robustly identifies cell types within the mouse cortex at sub-cortical layer resolution.

### *Plk2* expression correlates with vulnerability to cortical pSyn pathology in α-syn-tg mice

Given that tissue morphology is preserved throughout the IST workflow, we performed post-run IF with an antibody against α-synuclein phosphorylated at Ser129 in the same sections used for transcriptomics (Fig. 2a, Supplementary Fig. 1a). Images were co-registered with Xenium data, overlaid with Xenium-identified cells, and cells were assigned as pSyn^+^ or pSyn^-^ (Fig. 2a). We quantified vulnerability to pSyn pathology across cell types in the cortex as the percent of cells of each type which was pSyn^+^ (Fig. 2b). This analysis revealed that ExNs developed the vast majority of the pSyn pathology, whereas InNs and non-neuronal cells developed pathology infrequently (Fig. 2b). Within the ExN subtypes, however, we observed further selective vulnerability to pSyn pathology, particularly within L5; L5 ET neurons developed the most frequent pathology (∼50%), with L5 IT and L5 NP neurons developing much less pathology (∼30% and ∼20%, respectively) (Fig. 2b). L6 IT, L6 CT, and L6b neurons all displayed similar levels (∼40%), as did L2/3 and L4/5 IT neurons (∼35%) (Fig. 2b).

**Fig. 2.**
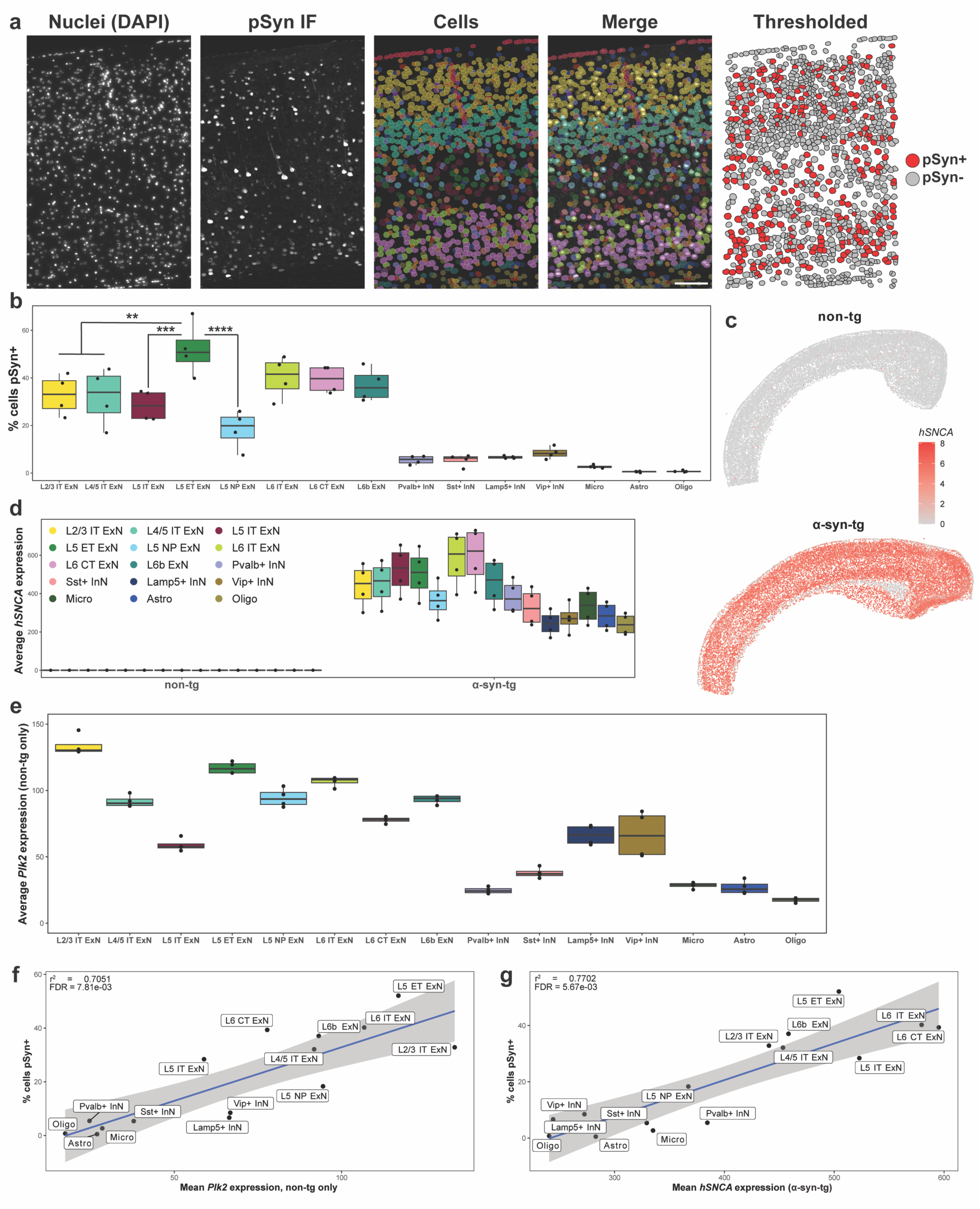
*Plk2* expression correlates with vulnerability to cortical pSyn pathology in α-syn-tg mice. **a** Representative cortical section from a female α-syn-tg brain (α-syn-tg #1) showing, from left to right, nuclei stained with DAPI imaged on the Xenium instrument, IF staining with an antibody against p-α-synuclein (clone EP1536Y) in the same sections used for Xenium, cells identified by Xenium via 5-μm nuclear expansion, pSyn IF and Xenium-defined cells overlaid, and the cells denoted as pSyn^+^ or pSyn^-^ following thresholding of the IF and assigning pSyn inclusions to individual cells (see Methods, “Xenium/pSyn overlap analysis”). Scale bar = 100 μm. **b** Quantification of vulnerability of selected cortical cell types to pSyn pathology in α-syn-tg mice (n = 4 α-syn-tg). Vulnerability of each cell type shown as percent of cells in each cluster which was identified as pSyn^+^, with each dot representing 1 sample. One-way ANOVA with Tukey’s multiple comparison’s test. Significance is shown relative to L5 ET ExNs, but full statistical results can be found in the Source Data file in our Zenodo repository (see Data Availability). **p < 0.01, ***p < 0.001, ****p < 0.0001. **c** Relative expression of *hSNCA* in a representative non-tg and α-syn-tg cortex. Specificity of hu-*SNCA* probes demonstrated with no cross-reactivity to ms-*Snca* in non-tg samples. **d** Quantification of *hSNCA* expression across cell types in the cortex of non-tg and α-syn-tg mice (n = 4 per genotype). Expression = expm1(log-normalized counts). **e** Quantification of *Plk2* expression across cell types in the cortex of non-tg mice (n = 4 non-tg). Expression = expm1(log-normalized counts). **f** Linear regression between average expression of *Plk2* by cell type across n = 4 non-tg samples and average percent of that same cell type which was pSyn^+^ across n = 4 α-syn-tg samples. **g** Linear regression between average expression of *hSNCA* by cell type and average percent of that same cell type which was pSyn^+^ across n = 4 α-syn-tg samples.

We next sought to understand the transcriptional underpinnings of the vulnerability and resilience of certain neuronal subtypes to pSyn pathology in this model. We first asked whether variable transgene expression across cell types may influence pathology rates, as pSyn pathology has been tied to α-synuclein expression in other systems^8, 23^. Since human and mouse α-synuclein sequences are highly conserved (∼95% protein sequence homology)^51^, we designed custom Xenium probes against human *SNCA* and mouse *Snca* which would not cross-react (Fig. 2c; custom probe sequences can be found in Supplementary Table 4). Visualization of *hSNCA* expression in non-tg and α-syn-tg brains confirmed specificity of the probes (Fig. 2c), as did subsequent quantification of *hSNCA* expression across cell types in non-tg and α-syn-tg samples (Fig. 2d). In both cases, non-tg samples show negligible *hSNCA* expression, whereas α-syn-tg samples demonstrate abundant expression across cell types (Fig. 2c, d).

As the *hSNCA* transgene is inserted on the X-chromosome of Line 61 α-syn-tg mice, sex differences are observed in the level of human α-syn protein and behavioral deficits in the model due to X-inactivation^47, 52^. Thus, we first sought to determine if we observed higher *hSNCA* expression in male α-syn-tg mice, and whether this corresponded with higher frequency of pSyn pathology (Supplementary Fig. 7). When we stratified *hSNCA* expression by sex, we observed that males displayed higher expression across every cell type in the cortex and hippocampus of the α-syn-tg mice (Supplementary Fig. 7a, c). Interestingly, however, this did not translate to higher pSyn pathology frequencies in male animals across those same cell types (Supplementary Fig. 7b, d). This relationship of human α-syn expression not generating more pSyn pathology in male α-syn-tg mice has been documented previously^52^, and needs to be investigated further. This does, however, suggest that combining males and females for further analysis of gene expression relationships with pSyn pathology across cell types is an appropriate strategy.

We observed variability when quantifying *hSNCA* expression across cortical cell types in α-syn-tg samples. Notably, ExNs displayed generally higher expression than InNs and non-neuronal cells (Fig. 2d). Within the ExN subtype, there was variability among the layer 5 and layer 6 subtypes, with L5 NP neurons expressing lower levels than L5 IT or L5 ET, and L6b neurons expressing lower levels than either L6 IT or L6 CT (Fig. 2d).

To test whether *hSNCA* expression correlated with pathology across cell types, we ran linear regressions between *hSNCA* expression and the percentage of cells pSyn^+^ for the different cell types in the cortex (Fig. 2g, Supplementary Table 5). Interestingly, we found that when performing this regression with all the major cell types in the cortex, *hSNCA* expression correlated with pSyn pathology (r^2^ = 0.7702, p < 0.0001, Fig. 2g, Supplementary Table 5). However, when we ran this same linear regression but only considering the ExN subtypes, *hSNCA* expression no longer correlated with pSyn frequency (r^2^ = 0.3635, p = 0.1136, Supplementary Table 5). Notable outliers in this analysis were L5 IT and L5 ET neurons, for which *hSNCA* expression was not predictive of the proportion of cells that showed pathology (Fig. 2b, d, g; Supplementary Table 5).

Given that *hSNCA* explained some, but not all the variance in pSyn pathology across cell types, we searched for additional genes that may influence pSyn pathology formation. We regressed the percent of cells in a cluster which were pSyn^+^ in α-syn-tg animals against gene expression from cells of the same cluster from non-tg animals (% cells pSyn^+^ ∼ non-tg GEx) (Supplementary Table 5). We chose this approach to avoid the likely outcome where, by regressing pSyn rates by cell type against gene expression from α-syn-tg animals, instead of observing upstream regulators of vulnerability to pathology, we would be observing downstream consequences of disease/pathology^15^. When performing this regression considering all the cell types in Fig. 2d, one of the most highly correlated genes was *Plk2*, the major kinase which phosphorylates α-synuclein at Ser129 in the central nervous system^53, 54^ (r^2^ = 0.7051, p < 0.0001, Fig. 2f). Across cell types, most ExN subtypes expressed relatively high levels of *Plk2* compared to InNs and non-neuronal cells (Fig. 2e), with the notable exception of L5 IT ExNs, which expressed *Plk2* at much lower levels than most other ExN subtypes, and at about half the levels of L5 ET ExNs (Fig. 2e). Interestingly, there were multiple genes whose expression correlated significantly with pSyn pathology in this analysis (Supplementary Table 5). However, these genes were either cell type marker genes (e.g., *Slc17a7*), expressed at low baseline levels (e.g., *Capn1*), or did not explain the large difference in pathology between layer 5 ExN subtypes (e.g., *Ext1, Hsph1*). Notably, when we performed this regression considering only the ExN subtypes, *Plk2* expression no longer correlated significantly with pathology rates (Supplementary Table 5), indicating that similarly to *hSNCA*, *Plk2* explains some, but not all the variance in pSyn pathology across cell types.

### Imaging spatial transcriptomics identifies major and rare hippocampal cell types

We next sought to analyze pSyn pathology by cell type in the hippocampus (Fig. 3a). Xenium identified 58,446 hippocampal cells across the 8 samples (Fig. 3b). Of note, we detected more cells in the α-syn-tg samples than the non-tg animals (Supplementary Fig. 8a). The sections from α-syn-tg animals were slightly more caudal than the non-tg sections, and thus the hippocampus was larger comparatively (Supplementary Fig. 9). All non-tg and α-syn-tg samples had similar numbers of detected transcripts per cell, unique genes per cell, and negative probe rates (Supplementary Fig. 8b-d).

**Fig. 3.**
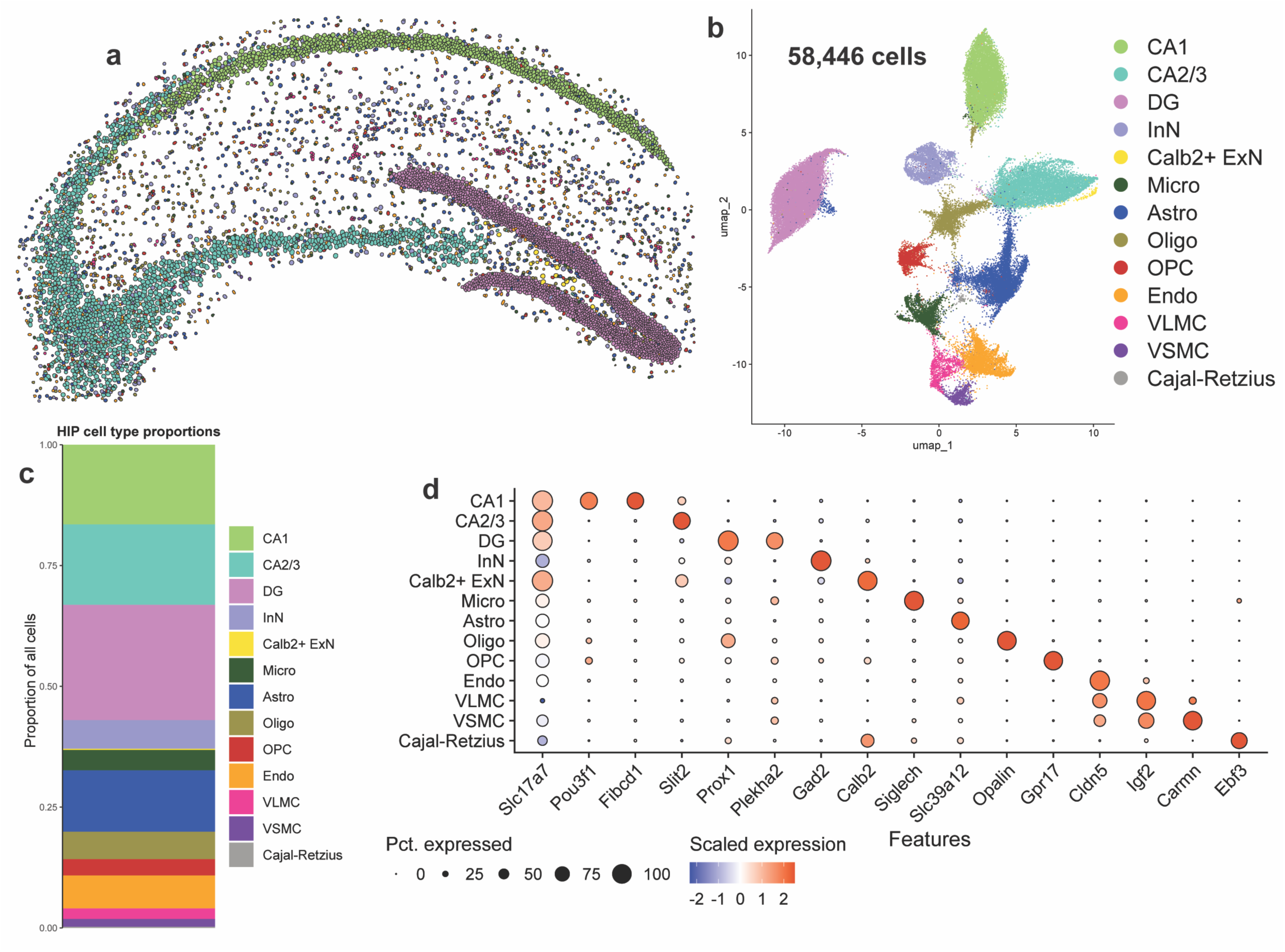
Imaging spatial transcriptomics identifies major and rare hippocampal cell types at single-cell resolution. **a** Hippocampal cells from one representative non-tg sample shown in physical space. Dots represent individual cells, colored by cell type. **b** All hippocampal cells from 8 samples (4 non-tg and 4 in α-syn-tg) shown in UMAP space (58,446 total cells). Dots represent individual cells, colored by cell type. **c** Proportions of all identified hippocampal cell types across all 8 samples. **d** Dotplot showing scaled expression of canonical marker genes for each hippocampal cell type.

Hippocampal cells were clustered into 13 transcriptionally distinct groups (Fig. 3b). We first visualized these cells in physical space, which revealed clear clustering of ExN subtypes by hippocampal sub-region, which were split into CA1, CA2/3, and dentate gyrus (DG) cells (Fig. 3a) and composed most of the cells identified in this brain region (Fig. 3c). These major ExN subtypes were transcriptionally distinct, with CA1 neurons expressing *Pou3f1* and *Fibcd1*, CA2/3 neurons expressing *Slit2*, and DG neurons expressing *Prox1* and *Plekha2* (Fig. 3c). Interestingly, Xenium identified a fourth ExN subtype marked specifically by the expression of *Calb2*, which encodes the Ca^2+^-binding protein calbindin 2 or calretinin (Fig. 3b, d). This cell type was extremely rare and spatially restricted, composing only ∼0.25% of all cells identified and being localized specifically to the hilus of the DG (Fig. 3a, c). Additionally, we identified 1 cluster of InNs, marked by *Gad2*, many of the same non-neuronal cells identified in the cortex (microglia, astrocytes, oligodendrocytes, OPCs, endothelial cells, VLMCs, and VSMCs), as well as a rare cluster of Cajal-Retzius cells marked by the expression of *Ebf3* (Fig. 3d).

### *Plk2* expression underlies vulnerability to hippocampal pSyn pathology in α-syn-tg mice

We next sought to determine which cell types were vulnerable to pSyn pathology in the hippocampus. By overlaying IF for pSyn with Xenium-identified cells, CA1 neurons had the highest proportion of pathology (∼35% of cells pSyn^+^), followed by CA2/3 neurons (∼15% of cells pSyn^+^); pSyn^+^ DG cells were very rare (Fig. 4a, b). As with the cortex, InNs and non-neuronal cells displayed a very low frequency of pathology (Fig. 4b).

**Fig. 4.**
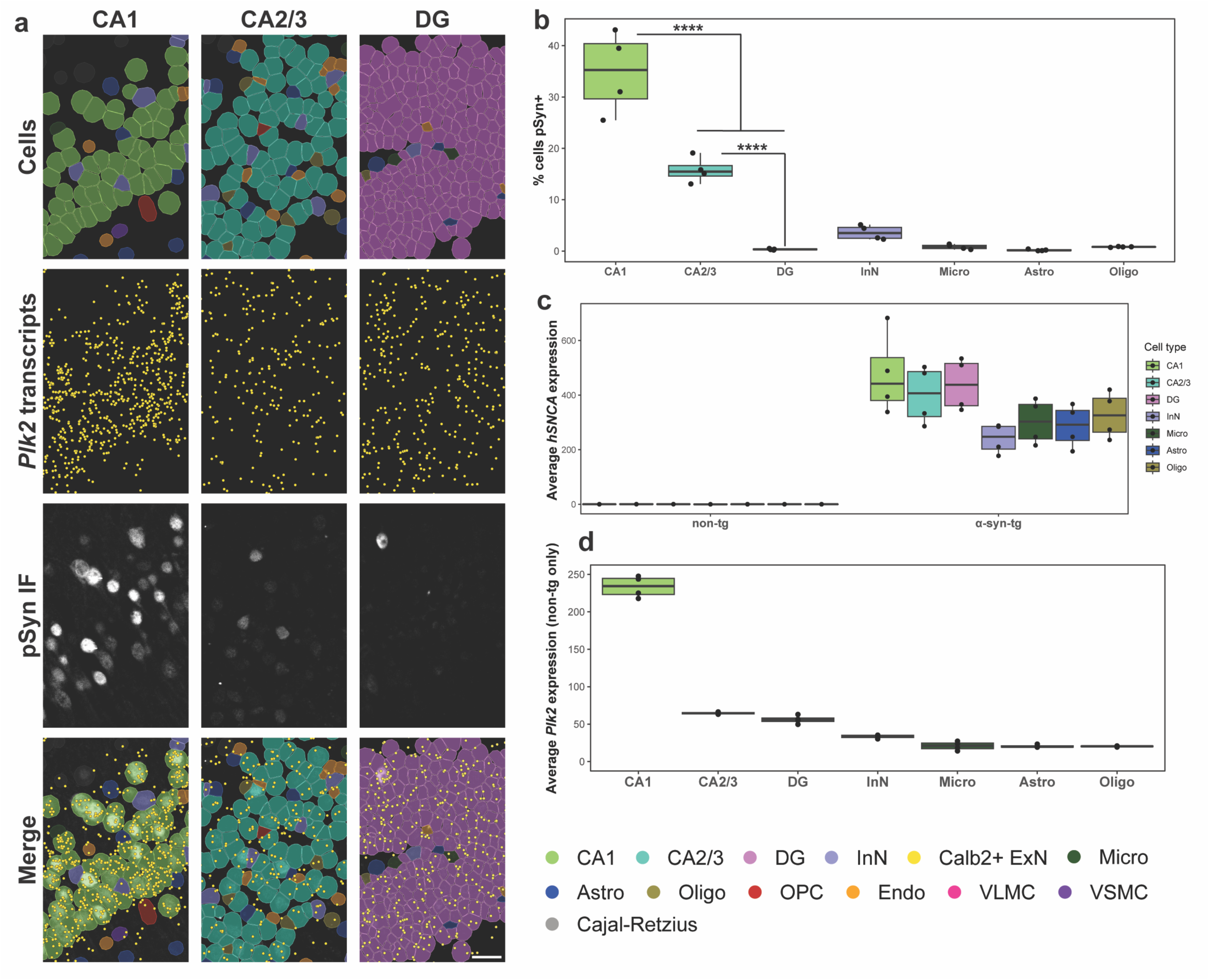
*Plk2* expression underlies vulnerability to hippocampal pSyn pathology in α-syn-tg mice. **a** Representative images from one α-syn-tg sample showing, from top to bottom, Xenium-defined cells, Xenium-detected *Plk2* transcripts (represented as individual points in yellow), IF staining with an antibody against p-α-synuclein (clone EP1536Y) in the same sections used for Xenium, and merged images. From left to right, images show the CA1, CA2/3, and DG hippocampal subfields. Scale bar = 30 μm. **b** Quantification of vulnerability of selected hippocampal cell types to pSyn pathology in α-syn-tg mice (n = 4 α-syn-tg). Vulnerability of each cell type shown as percent of cells in each cluster which was identified as pSyn^+^, with each dot representing 1 sample. One-way ANOVA with Tukey’s multiple comparison’s test. Significance is shown between major excitatory neuron subtypes, but full statistical results can be found in the Source Data file in our Zenodo repository (see Data Availability). ****p < 0.0001. **c** Quantification of *hSNCA* expression across cell types in the hippocampus of non-tg and α-syn-tg mice (n = 4 non-tg and 4 α-syn-tg). Expression = expm1(log-normalized counts). **d** Quantification of *Plk2* expression across cell types in the hippocampus of non-tg mice (n = 4 non-tg). Expression = expm1(log-normalized counts).

Given such striking differences in vulnerability to pSyn pathology, we sought to investigate the transcriptional underpinnings of these discrepancies. We first quantified the average *hSNCA* expression for each cell type in each sample (Fig. 4c), which revealed that on average, CA1, CA2/3, and DG neurons all expressed relatively even *hSNCA* levels, while InNs and all non-neuronal cells expressed slightly lower levels (Fig. 4c). Given that the three major ExN subtypes expressed similar levels of *hSNCA*, but nonetheless had dramatically different rates of pSyn pathology, we again turned to other genes which could potentially explain this effect.

When we quantified the expression of *Plk2* across the hippocampal cell types, we found that expression was ∼4-fold higher in CA1 neurons than in either CA2/3 or DG neurons (Fig. 4d), likely explaining the relatively much higher rates of pSyn pathology in CA1 neurons in this model. Again, however, *Plk2* did not seem to explain all of the vulnerability to pSyn pathology, as the difference in vulnerability between CA2/3 and DG neurons (∼15% vs. ∼0%) is not explainable by *hSNCA* or *Plk2* expression (Fig. 4b-d). This difference could potentially be explained other genes not measured here, or alternatively through Plk2 at the protein level, either expression or kinase activity.

### *Plk2* may explain differences in pSyn vulnerability within some cell types

While the above data demonstrate that differences between cell types in propensity to pSyn pathology relates to expression of the kinase:substrate pair, we note that there is also likely to be within-cluster vulnerability. For example, even in the most vulnerable cell type, L5 ET neurons, only ∼50% of cells developed pathology (Fig. 2b). We therefore analyzed our IST data to nominate genes that might explain variability in pathology within specified cell types.

We first turned to hSNCA expression, which we quantified on a per-cell basis (Fig. 5a). We found that, on average, pSyn^+^ cells of every ExN subtype in the cortex expressed slightly higher *hSNCA* levels than pSyn^-^ cells (Fig. 5a), again indicating that α-synuclein expression is a contributing factor to pathology formation. Interestingly, however, there are many neurons which expressed high *hSNCA* but were pSyn^-^, and cells that expressed low *hSNCA* yet were pSyn^+^ (Fig. 5a, b), again indicating that *hSNCA* expression is not the sole determinant of vulnerability to pathology. We therefore subsetted cells from each ExN subtype which were either pSyn^-^ and above the 75^th^ percentile by *hSNCA* expression (pSyn^-^ / *hSNCA*-high) or cells which were pSyn^+^ and below the 25^th^ percentile by *hSNCA* expression (pSyn^+^ / *hSNCA*-low) (Fig. 5a, b). This subsetting resulted in groups of cells from each ExN subtype where the pSyn^+^ cells had lower *hSNCA* expression than the pSyn^-^ cells, on average (Fig. 5a, b). We then performed pseudo-bulk differential expression testing on these groups of cells, using pSyn status as the variable of interest (Fig. 5c). Plotting differentially expressed genes (DEGs) between these two groups across neuronal subtypes confirmed that in these comparisons, pSyn^+^ cells expressed significantly lower *hSNCA* than the pSyn^-^ cells (Fig. 5c, all significant results in Supplementary Table 6).

**Fig. 5.**
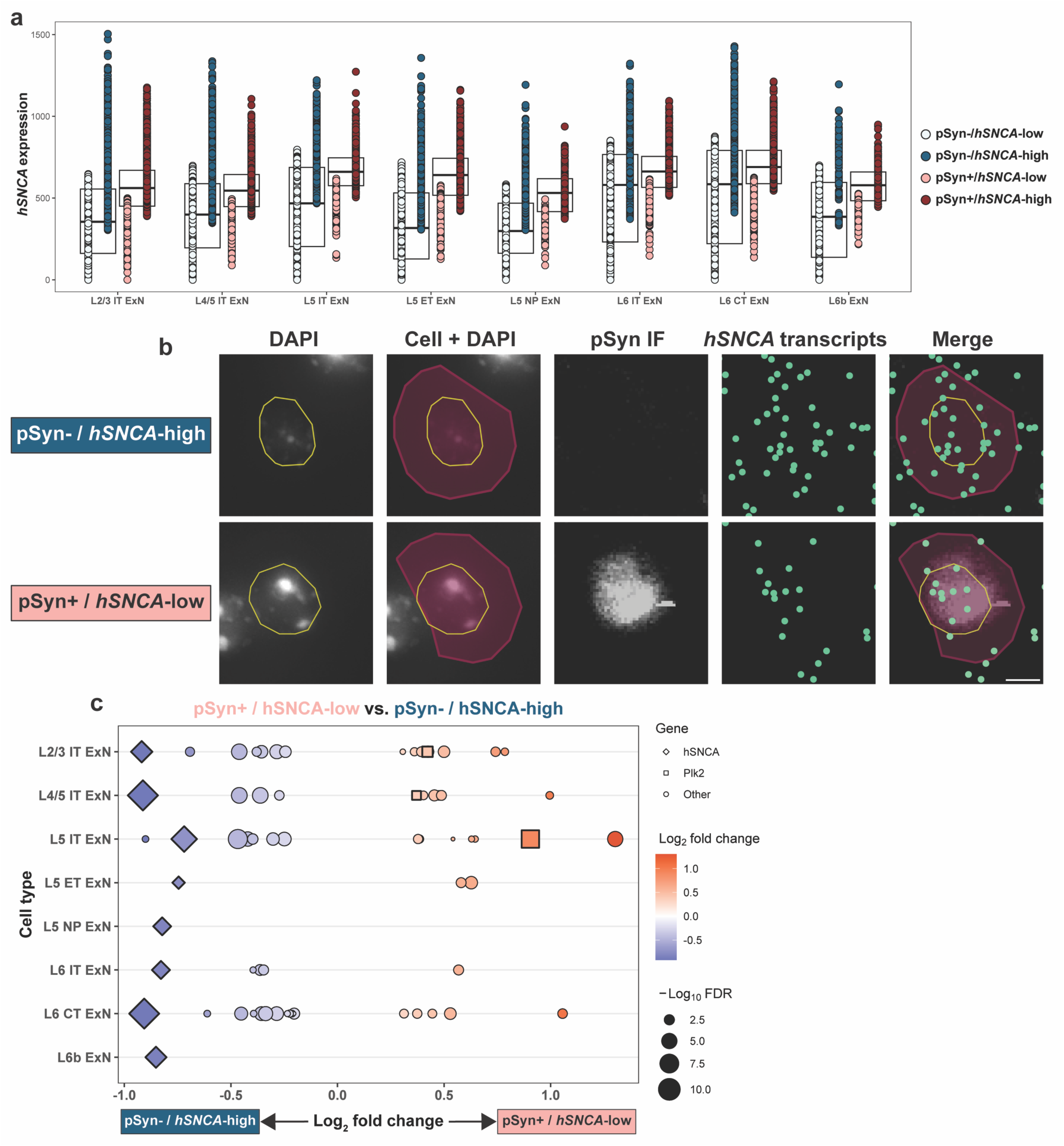
*hSNCA* and *Plk2* expression are implicated in within-cluster pSyn vulnerability. **a** *hSNCA* expression in all general ExN subtypes. Each dot represents an individual cell. Cells are colored according to their pSyn status (blue = pSyn^-^, red = pSyn^+^), and their *hSNCA* expression, where dark blue indicates cells in the top 25% by *hSNCA* expression (pSyn^-^ / *hSNCA*-high) and light red indicates pSyn^+^ cells in the bottom 25% by *hSNCA* expression (pSyn^+^ / *hSNCA*-low). These pSyn^-^ / *hSNCA*-high and pSyn^+^ / *hSNCA*-low cells were used for differential expression analysis in **(c)**. Expression = expm1(log-normalized counts). **b** Representative images of single L5 IT neurons which were either pSyn^-^ / *hSNCA*-high (top) or pSyn^+^ / *hSNCA*-low (bottom). Images were taken from the same animal (α-syn-tg #1). From left to right, images show nuclei stained with DAPI imaged on the Xenium instrument outlined in yellow, Xenium-identified cells overlaid with their nuclei, IF staining against p-α-synuclein, Xenium-detected *hSNCA* transcripts (represented as individual points in green), and pSyn IF, *hSNCA* transcripts, nuclei outlines, and cells overlaid. Scale bar = 5 μm. **c** DEGs for pseudo-bulk differential expression analysis performed on the cells shown **(a)**, directly comparing pSyn^+^ / *hSNCA*-low and pSyn^-^ / *hSNCA*-high cells. All genes shown are significant DEGs (FDR < 0.05); enriched DEGs are shown in red, with depleted DEGs shown in blue. Size of the shape represents the -log_10_FDR. *hSNCA* is shown as a diamond, *Plk2* is shown as a square, and other genes are represented as circles.

We then sought to understand if any of the DEGs could help explain why the *hSNCA*-high cells did not develop pSyn pathology while the *hSNCA*-low cells did. *Plk2* again emerged as a potential explanation for this phenomenon, being enriched in pSyn^+^ / *hSNCA*-low cells of three subtypes (L2/3, L4/5, L5 IT ExNs) (Fig. 5c). Interestingly, L5 IT ExNs, which had the lowest baseline *Plk2* expression of any ExN subtype (Fig. 2e), had the largest difference in *Plk2* expression between pSyn^+^ / *hSNCA*-low and pSyn^-^ / *hSNCA*-high cells (Fig. 5c), indicating that a subpopulation of L5 IT ExNs which is *Plk2*-high compared to the rest of the cluster are the cells which develop pSyn pathology.

In addition, we performed differential expression analysis between pSyn^+^ and pSyn^-^ cells from the hippocampus (Supplementary Table 7). Due to the very low number of pSyn^+^ CA2/3 and DG cells, we could not reliably perform the same percentile-based analysis as was done for the cortex. Interestingly, while there were not many significant DEGs in this analysis, we do note that pSyn^+^ DG cells expressed significantly higher *Plk2* than pSyn^-^ DG cells (log_2_FC = 1.24), again pointing to *Plk2* as a determining factor in pSyn pathology formation within certain cell types (Supplementary Table 7).

This data again implicates the kinase:substrate relationship of *Plk2* and α-synuclein, now also in determining which cells within certain neuronal subtypes are vulnerable to pSyn pathology. This also potentially suggests a reciprocal relationship of expression of these two genes, where cells with lower α-synuclein expression may still develop pSyn pathology if they express higher *Plk2* levels, and *vice versa*.

### Imaging spatial transcriptomics reveals conserved transcriptional dysfunction downstream of *hSNCA* overexpression and pSyn pathology

Finally, we aimed to determine downstream transcriptional effects of *hSNCA* overexpression and the presence of pSyn pathology in α-syn-tg neurons. We performed two separate analyses across the general ExN subtypes from the cortex (Fig. 6a). First, we performed pseudo-bulk differential expression analysis with three separate comparisons, comparing non-tg cells to both pSyn^+^ and pSyn^-^ α-syn-tg cells (Fig. 6b, c), and comparing pSyn^+^ and pSyn^-^ α-syn-tg cells to each other (Fig. 6d). This analysis was designed to determine the downstream transcriptional effects of *hSNCA* expression and the presence of pSyn pathology separately. We then performed a generalized linear model (GLM) analysis, regressing the expression of genes found to be differentially expressed via pseudo-bulk analysis against *hSNCA* expression (GEx ∼*hSNCA*) in individual cells (Fig. 6e). This analysis was performed only on α-syn-tg cells, and given that pSyn^+^ cells had higher average *hSNCA* expression than pSyn^-^ cells, intended to help disentangle those differentially expressed genes (DEGs) which were due directly to *hSNCA* overexpression versus those which were a result of pSyn pathology.

**Fig. 6.**
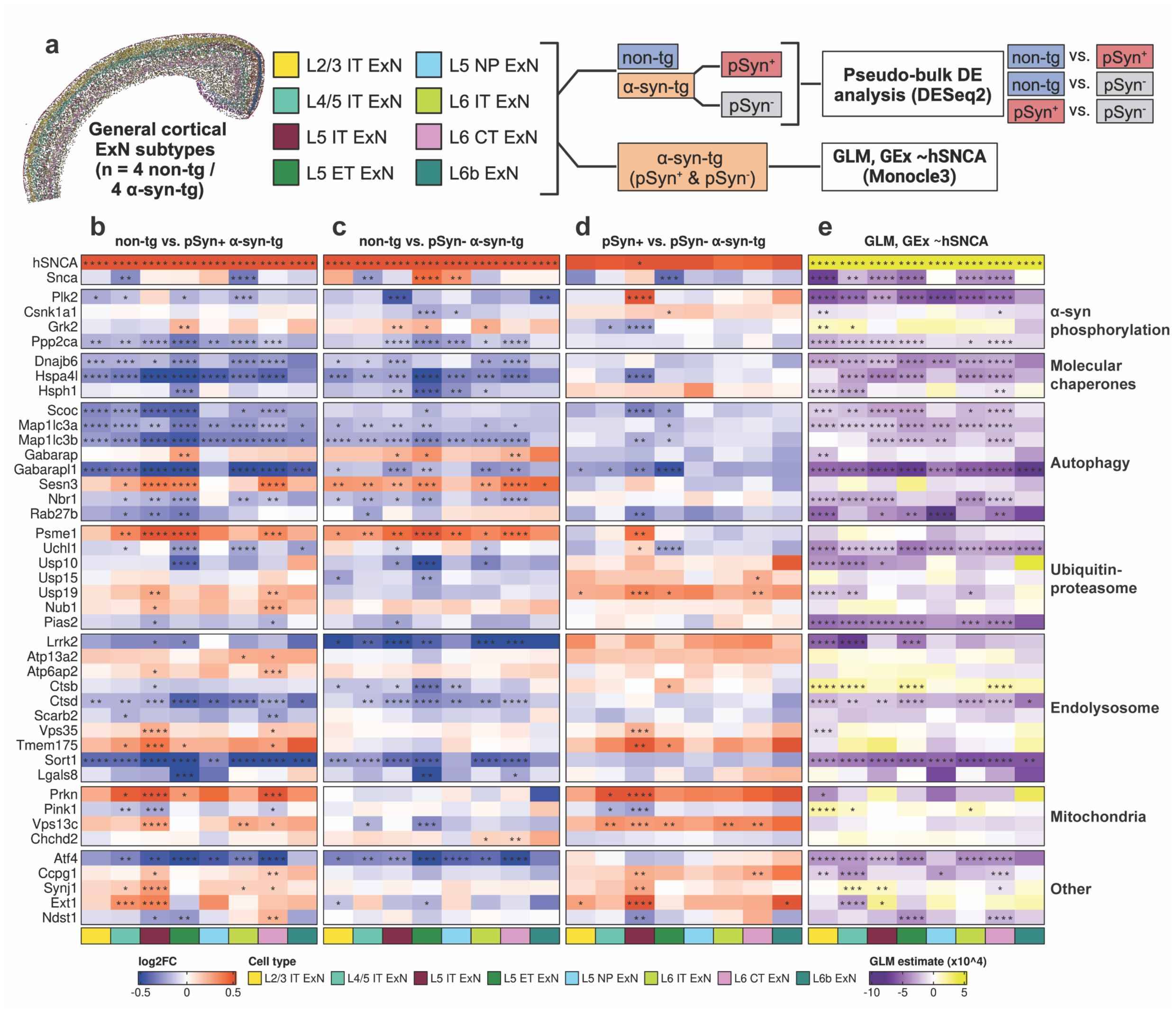
Differential expression analysis of pSyn^+^ and pSyn^-^ cortical ExNs reveals conserved dysfunction in α-syn-tg neurons. **a** General ExN subtypes were split into 3 categories (non-tg, pSyn^+^ α-syn-tg, and pSyn^-^ α-syn-tg). Pseudo-bulk differential expression analysis was run using DESeq2, with 3 comparisons: non-tg vs. pSyn^+^ α-syn-tg, non-tg vs. pSyn^-^ α-syn-tg, and pSyn^+^ vs. pSyn^-^ α-syn-tg. Next, Monocle3 was used to perform a generalized linear model (GLM) analysis on only the α-syn-tg cells, independent of pSyn status, regressing the expression of the pseudo-bulk-identified DEGs (GEx) against the expression of *hSNCA* in single α-syn-tg cells, to determine if expression of the DEGs correlated with *hSNCA* expression. Created in BioRender. https://BioRender.com/d72l899. **b-d** Heatmap of differentially expressed genes for non-tg vs. pSyn^+^ α-syn-tg cells (**b)**, non-tg vs. pSyn^-^ α-syn-tg cells **(c)**, and pSyn^+^ vs. pSyn^-^ α-syn-tg cells (**d)**. Each column represents an ExN subtype (see legend). *FDR < 0.05, **FDR < 0.01, ***FDR < 0.001, ****FDR < 0.0001. **e** Heatmap of results of Monocle3 GLM analysis of DEG gene expression vs. hSNCA expression at the single-cell level in α-syn-tg cell (GEx ∼*hSNCA*). Color of the cell represents the correlation direction and strength (purple = negative correlation, yellow = positive correlation). Each column represents an ExN subtype (see legend). *q < 0.05, **q < 0.01, ***q < 0.001, ****q < 0.0001. Full differential expression results and GLM results can be found in Supplementary Table 8.

We first examined the abundance of DEGs across the ExN subtypes for each of the different comparisons (Supplementary Fig. 10). We observed high overlap between the non-tg vs. pSyn^-^ and non-tg vs. pSyn^+^ α-syn-tg comparisons for most cell types (Supplementary Fig. 7a). However, we note that each cell type exhibited DEGs which were either pSyn-specific or pSyn-independent when compared to non-tg (Supplementary Fig. 10).

We hypothesized that the transcriptional response in α-syn-tg neurons may be conserved across subtypes. Therefore, we only considered DEGs which were significantly up- or downregulated in two or more cell types in the same direction (full lists of DEGs for all comparisons, along with GLM results, can be found in Supplementary Table 8). We manually split the DEGs fitting those criteria based on their known function pertaining to α-synuclein homeostasis; these groupings were determined prior to designing the custom Xenium panel, based on the literature.

We first observed lower expression of molecular chaperones (*Dnajb6*, *Hspa4l*, *Hsph1*), and autophagy-related genes (*Scoc*, *Map1lc3a*, *Map1lc3b*, *Gabarapl1*, *Nbr1*, *Rab27b*) in α-syn-tg neurons, and expression levels of the same genes were negatively correlated with higher levels of *hSNCA* expression (Fig. 6b-e). There were some exceptions (*Gabarap*, *Sesn3*) which were more highly expressed in α-syn-tg neurons, and expression of these genes did not correlate with hSNCA expression (Fig. 6b-e).

We also observed lower expression of *Plk2* in α-syn-tg neurons (Fig. 6b-d), and *Plk2* expression negatively correlated with *hSNCA* expression across most cell types (Fig. 6e). Along with *Plk2*, we also observed lower expression of *Ppp2ca*, encoding a subunit of Pp2a, a Ser/Thr phosphatase that dephosphorylates α-synuclein at Ser129^55^ (Fig. 6b-d).

Genes related to proteolysis, notably the ubiquitin-proteasome system, showed mixed direction of effect, with some genes (*Psme1*, *Nub1*) showing higher expression in α-syn-tg neurons, while others (*Uchl1*, *Usp10*) showed lower expression (Fig. 6b-d). Of these genes, *Usp19* showed a pSyn-dependent response, as its expression was increased in pSyn^+^ α-syn-tg neurons compared to non-tg but was not altered in pSyn^-^ neurons (Fig. 6b-e).

We also noted several genes related to endolysosomal function that showed differential regulation in this analysis. For example, the lysosomal protease *Ctsd* was downregulated in α-syn-tg neurons, along with endosomal sorting protein *Sort1* (Fig. 6b-d); both genes showed a negative correlation between expression and *hSNCA* expression (Fig. 6e). Interestingly, *Ctsd* showed consistent downregulation across pSyn-cell types, but was not downregulated in most pSyn^+^ cells, showing a positive correlation with *hSNCA* expression (Fig. 6b-e). We also noted upregulation of several endolysosomal genes (*Atp13a2*, *Atp6ap2*, *Vps35*, *Tmem175*) that were specific to pSyn^+^ α-syn-tg neurons (Fig. 6b-d). Importantly, expression of these genes did not correlate with *hSNCA* expression, indicating that their upregulation in pSyn^+^ cells is likely due to the presence of pSyn pathology rather than higher *hSNCA* expression in those cells (Fig. 6e).

Finally, we noted that of four mitochondrial DEGs, *Prkn* was upregulated in pSyn^+^ neurons while exhibiting no change in pSyn^-^ cells (Fig. 6b-d). Similarly, *Vps13c* was upregulated in pSyn^+^ cells while being downregulated in some pSyn^-^ subtypes (Fig. 6b-d). In contrast, *Pink1* was downregulated in pSyn^+^ cells while not being differentially expressed in pSyn^-^ cells, and *Chchd2* was upregulated in pSyn^-^ cells while showing no change in pSyn^+^ cells (Fig. 6b-d). Accordingly, expression of these genes did not show strong or conserved correlation in either direction when regressed against *hSNCA* expression (Fig. 6e), indicating that differential expression of these mitochondrial genes is likely also due to the presence of pSyn pathology rather than *hSNCA* expression.

## Discussion

The major goals of this study were three-fold: first, to identify neuronal subtypes vulnerable and resistant to phospho-α-synuclein pathology in a transgenic mouse model of human α-synuclein overexpression; second, to investigate the transcriptional underpinnings of that vulnerability; and third, to determine downstream transcriptional effects of *hSNCA* overexpression and the presence of α-synuclein pathology.

Prior studies have examined cellular vulnerability to α-synuclein pathology in PD/DLB and in the α-synuclein preformed fibril model^5, 22^. However, to our knowledge, no study has examined this in a transgenic synucleinopathy model. Given the widespread use of transgenic models in the PD/DLB and Alzheimer’s disease (AD) fields, it is important to compare the consistency of pathology distribution between animal models and their accuracy when compared to human disease.

We first evaluated vulnerability to pSyn pathology by cell type in the cortex and hippocampus of α-syn-tg mice. In the cortex, we found that ExNs were broadly vulnerable, while InNs did not develop pathology, despite overexpressing *hSNCA*. We also found further selective vulnerability among the ExN subtypes, determining that L5 ET neurons developed the most frequent pathology, while other subtypes even in the same cortical layer (L5 IT and L5 NP) developed pathology less frequently. Outer-layer (L2/3 IT and L4/5 IT) and deep-layer (L6 IT, L6 CT, and L6b) neurons all developed pathology at similar frequencies. Interestingly, we note that the most severe pathology, which extended to the cytoplasm and axons, was also most prevalent in L5 ET neurons, while other neuronal subtypes mainly had pathology restricted to the nucleus. In the hippocampus, we found that InNs were broadly resistant to pSyn pathology, and among the major ExN subtypes, CA1 neurons developed pathology approximately twice as frequently as CA2/3 neurons, while DG neurons exhibited almost no pathology.

Recent work performed in human tissue and the α-synuclein PFF model sheds further light on selective neuronal vulnerability to pSyn pathology. Specifically, it has been demonstrated that in the cortex and amygdala of mice injected with α-synuclein PFFs, ExNs are broadly vulnerable to pSyn pathology, while InNs are resistant ^5, 22^, consistent with our observations in the α-syn-tg mouse model. However, when specifically comparing among ExN subtypes in the cortex of PFF mice and PD cases, L5 IT neurons were vulnerable to pSyn pathology, while L5 ET neurons were resilient^5^. It is possible that different model systems (i.e., transgenic vs. fibril seeding) may engage different mechanisms of vulnerability in different cell types.

In the hippocampus, work in α-synuclein PFF-injected mice and human DLB tissue has identified the CA2/3 subfield as the most vulnerable to pSyn pathology and subsequent degeneration^8, 20, 56^. This contrasts with our current findings in the α-syn-tg model, in which we found CA1 to have the most pSyn pathology. Interestingly, however, prior work in the same α-syn-tg model showed that CA3 neurons, but not CA1 neurons, were vulnerable to degeneration, which was attributed to higher expression of metabotropic glutamate receptor 5 (mGluR5) in the vulnerable cells^57^. These results imply that there are distinct mechanisms of vulnerability to pSyn pathology compared to degeneration, and that pSyn pathology may not necessarily precede degeneration of affected neurons. In support of this possibility, it has been demonstrated in a transgenic mouse model of tauopathy that neurons bearing neurofibrillary tangles (NFTs) were less vulnerable to degeneration than NFT-free neurons^58^. Further work is needed in mouse models and human tissue of synucleinopathy and other proteinopathies to determine if pathology is invariably upstream of cell death or if separate mechanisms govern vulnerability to these distinct events, and if this process occurs in a cell type-dependent manner.

We next investigated which genes included in our Xenium panel may be responsible for differences in pSyn vulnerability across and within cell types. In the context of synucleinopathies, prior work has implicated endogenous α-synuclein expression in the selective vulnerability of certain neuronal populations, at the regional and subtype level^8, 24, 59^. For example, ExNs, which are broadly vulnerable to pathology, express high levels of α-synuclein, where InNs, which are resilient to pathology, express low levels of α-synuclein^24^. Additionally, the CA2/3 region of the hippocampus expresses the highest levels of endogenous α-synuclein and is accordingly the most vulnerable to pathology in the PFF model^8^. However, α-synuclein expression does not seem to account for the whole picture of vulnerability to pathology; for example, DA neurons of the VTA express similarly high α-synuclein levels as those in the neighboring SNc, and yet do not develop pathology or degenerate in PD^24^. Work in the context of tauopathy found that certain protein clearance pathways, centered around the cochaperone BAG3, are expressed more highly in resilient InNs compared to vulnerable ExNs under baseline conditions, potentially implicating deficient clearance leading to protein aggregation and pathology in vulnerable cell types^29^. We thus hypothesized that similar cell-intrinsic differences might influence pSyn pathology formation in our synucleinopathy model, explaining differences in vulnerability between cell types which express similar levels of α-synuclein.

Our results implicate expression of the substrate:kinase pair of α-synuclein and Plk2 in the development of pSyn pathology in this model. In both cortex and hippocampus, ExNs showed higher *hSNCA* expression than InNs and non-neuronal cells, likely contributing to the absence of pathology in the latter cell types. However, our data supports the hypothesis that α-synuclein expression alone does not fully explain vulnerability to pathology. This observation is especially evident in the hippocampus, where CA1, CA2/3 and DG neurons expressed very similar *hSNCA* levels yet had highly variant pathology rates. Similarly, in the cortex, *hSNCA* expression does not correlate with pathology rates across cell types when only analyzing the ExN subtypes, particularly evident when comparing L5 IT and L5 ET neurons, which have nearly identical *hSNCA* expression but greatly different pathology rates.

In addition to *hSNCA*, *Plk2* expression correlates well with pathology rates across all cell types. Notably, InNs and non-neuronal cells generally displayed much lower *Plk2* expression than ExNs. Specifically within cortical layer 5, *Plk2* expression is around twice as high in vulnerable L5 ET neurons compared to the more resilient L5 IT neurons, likely explaining the discrepancy in pSyn pathology despite similar *hSNCA* expression between these two populations. In the hippocampus, this relationship becomes more evident. CA1 neurons express around four-fold higher *Plk2* than CA2/3 or DG neurons, correlating with their much higher vulnerability. However again, like with *hSNCA*, *Plk2* expression alone seemingly does not fully explain the vulnerability, as, for example, the difference in pathology rates between CA2/3 and DG neurons cannot be explained by differences in *Plk2* expression. It is likely that a complex interplay of multiple transcriptional, structural, and functional factors dictates a cell type’s vulnerability to pathology, and further work will be required to uncover more of these contributors.

Plk2 has been characterized as the primary kinase which phosphorylates α-synuclein at Ser129 *in vitro* and in the central nervous system^53, 54^. Our study supports and expands upon this work, implicating *Plk2* expression in the vulnerability of certain cell types to developing pSyn pathology in the α-syn-tg mouse model. However, the exact role of phosphorylated α-synuclein in both the physiological and disease states has remained elusive, and whether preventing α-synuclein phosphorylation by inhibiting Plk2 would be beneficial to slow disease progression is unclear. For example, increased α-synuclein phosphorylation induced by Plk2 overexpression in dopaminergic neurons was not neurotoxic^60^, and in fact, phosphorylation by Plk2 has been shown to promote the autophagic clearance of α-synuclein and reduce cytotoxicity^61^. Multiple studies in the α-synuclein PFF model have also shown that Plk2 inhibition or genetic deletion reduces phosphorylation of physiological, but not aggregated α-synuclein^62, 63^. Nevertheless, the abundance of pSer129 α-synuclein in the disease state compared to health^3, 4^ implies importance of this post-translational modification in disease. Thus, more work is needed to uncover the precise mechanisms and function of α-synuclein phosphorylation in health and disease, and accordingly whether Plk2 inhibition is a viable therapeutic target in synucleinopathies.

We also performed differential expression analysis to examine the transcriptional effects of *hSNCA* overexpression and pSyn pathology. We take four major conclusions from this data. First, we observed gene expression changes across multiple genes split into functionally relevant groups (e.g., autophagy), potentially indicative of pathway-level dysfunction. Second, many of these transcriptional changes were conserved across cell types, indicating that the neuronal response to *hSNCA* overexpression and pSyn pathology is not subtype-specific. Third, while many transcriptional changes occurred in the same direction in both pSyn^-^ and pSyn^+^ cells, indicative of a general response to *hSNCA* overexpression, there were also several genes, particularly enriched in endolysosomal and mitochondrial pathways, which were only altered (or changed in the opposite direction) in pSyn^+^ cells, indicative of a pSyn-specific transcriptional response. Fourth, many DEGs were also PD risk genes, potentially indicative of convergence of molecular pathways involving these risk genes around α-synuclein.

Given the use of our targeted panel, we do not expect to capture the full range of the cellular responses to α-synuclein. However, we believe that our panel was designed to be sensitive to many of the pathways involved in α-synuclein pathology, particularly focused on intracellular mechanisms of α-synuclein aggregation, clearance, and toxicity. Notably, our panel did not cover genes involved in inflammation, which is known to play a role in PD pathogenesis and in the α-syn-tg mouse model^64, 65^, so as higher-plex panels emerge, this will be an area of interest, particularly with regards to the spatiotemporal evolution of inflammation in synucleinopathies.

We observed transcriptional dysregulation of pathways central to α-synuclein homeostasis in neurons, including molecular chaperones, autophagy, the ubiquitin-proteasome and endolysosomal systems, and mitochondria. Each of these systems play key roles in α-synuclein processing, and each has evidence of potentially causal dysregulation in synucleinopathy^66–71^. There were many genes across almost all groups which were either enriched or depleted in both pSyn^+^ and pSyn^-^ neurons compared to non-tg, and which showed strong, non-cell type-specific correlation with *hSNCA* expression (e.g., *Scoc*, *Map1lc3b*, *Gabarapl1*, *Uchl1*, *Ctsd*, *Sort1*), indicative of a general response to proteostatic stress induced by human α-synuclein overexpression. However, there were also many genes whose expression was specifically altered in pSyn^+^ neurons and where expression did not correlate with *hSNCA* (e.g., *Pink1*, *Prkn*, *Vps35*), indicating a more specific cellular response to phosphorylated α-synuclein. Interestingly, this has been demonstrated previously, where a truncated form of α-synuclein phosphorylated at Ser129, termed pSyn*, was identified in human PD brains and in α-synuclein PFF-treated neurons and mice and was highly mitotoxic, inducing mitochondrial damage and mitophagy in the neurons^72^. Finally, many of the transcriptional changes which we observed, pSyn pathology-dependent or not, have been reported in other models of synucleinopathy or in human tissue (e.g., *Dnajb6*, *Hspa4l*, *Hsph1*, *Gabarapl1*, *Map1lc3b*, *Ctsb*, *Ctsd*, *Pink1*, among others)^5, 73–76^, indicating that broader α-synuclein-related cellular dysfunction is reflected in α-syn-tg mice.

Importantly, our data suggests that while α-syn-tg neurons with pSyn pathology show unique signatures of transcriptional dysfunction compared to neighboring pSyn^-^ α-syn-tg cells, those pSyn^-^ cells still experience significant alterations compared to healthy control cells. While this may be expected in a transgenic model where most cells are being impacted by transgene expression, this statement likely holds true in other models (i.e., α-synuclein PFF) and in human disease, as it does for AT8 tau-negative cells from AD brains compared to healthy controls^28^.

Given the abundance of synaptic α-synuclein pathology in the PD/DLB cortex^77^, for example, it is entirely likely that neurons which may not harbor LBs are still dysfunctional, transcriptionally and functionally. Thus, simply comparing transcriptomes of pSyn^+^ neurons to those of surrounding pSyn^-^ neurons within the same samples may not capture the full range of dysfunction in both populations of cells when compared to the healthy state. This necessitates independent comparisons of pSyn^+^ and pSyn^-^ diseased cells to cells from healthy controls in addition to within-sample comparisons of pSyn^+^ and pSyn^-^ cells.

There are several limitations to the current study. First, imaging spatial transcriptomics is a newer technology, and bioinformatic analysis techniques are still under development. Specifically, it is likely that instrument hardware and software (i.e., imaging quality, optical crowding, processing speeds), platform chemistries (i.e., probe set sizes, multiplexed protein targets for segmentation), and analysis techniques (i.e., segmentation algorithms, gene count normalization, differential expression) will evolve quickly over the coming years as benchmarking studies are conducted and new pipelines are established specifically for analyzing IST data. As such, we hope that this study and its associated raw data will serve as a resource for the community in developing these tools.

In addition, the mouse model we chose to use here, Line 61, has several unique features that may limit generalizability to other models. We chose this model mainly due to the large amounts of pSyn pathology it develops, which adequately powered our differential expression between pSyn^+^ and pSyn^-^ cells. However, transgene expression is driven under the *Thy1* promoter inserted on the X-chromosome, leading to sex effects which we confirm here. It is also possible that transgene expression, and by extension, pathology formation, is driven by promoter copy number or endogenous expression of *Thy1* transcription factors. α-Synuclein overexpression as a model also has limitations, in that the many-fold increase in protein levels^31^ may induce non-PD-relevant neuronal stress, and it is unclear whether phenotypes observed in transgenic and AAV-based α-synuclein overexpression models are due to overexpression or the formation of disease-relevant pathological α-synuclein assemblies^78^. However, our observation of transcriptional changes in α-syn-tg neurons which were reflective of alterations seen in other animal models of PD and in LB-bearing neurons of the human brain indicate that we are observing disease-relevant changes, to the extent that is possible in an overexpression model. Finally, this mouse line does not exhibit *bona fide* Lewy pathology; rather, neurons develop nuclear and cell body α-syn inclusions, some of which are proteinase-K resistant, as demonstrated here. Furthermore, however, no synucleinopathy mouse model develops true Lewy pathology, the closest currently being the α-syn preformed fibril model. Now that technology exists to extensively the profile molecular identities of cells with co-detection of pathology in the same sections, we believe it is important to re-evaluate animal models which have been in use for decades, to determine how faithfully each recapitulates aspects of human Lewy body disease (i.e., which cell types develop pathology and the downstream impacts of that pathology), which is what we sought to do in this study with the Line 61 model.

Overall, imaging spatial transcriptomics represents a promising avenue for neuropathological research. The ability to combine RNA, protein, and other -omic data in the same tissue sections while maintaining spatial fidelity is invaluable, allowing for the multimodal integration of high-dimensional data with higher confidence than when using any set of individual methods in parallel (i.e., analyzing serial sections or sets of isolated cells). Here, we utilized this novel technology in a mouse model of α-synucleinopathy, identifying neuronal subtypes which were vulnerable to pathology in the model, determining why they were vulnerable, and profiling downstream effects of pathology. Our study provides not only new biological insights into the molecular context of α-synuclein pathology, but also provides a framework for future neuropathological studies to expand upon, utilizing a similar technical toolkit to answer similar questions in other disease models and contexts.

## Acknowledgements

We appreciate Dr. Zu-Xi Yu at the Pathology Core in the National Heart, Lung, and Blood Institute (NHLBI) for allowing us to use the paraffin processor and embedding center. We also thank Dr. Xylena Reed at the Center for Alzheimer’s and Related Disorders (CARD) for facilitating the Xenium run, Dr. Jinhui Ding for assisting with the data transfer to GEO, Dr. Eliezer Masliah for providing the Line 61 mice and for intellectual input regarding the line, and members of the Cell Biology and Gene Expression Section for intellectual discussion and input.

## Author contributions

Conceptualization: LH-P, MI, MRC; Data acquisition: LH-P, MI; Data analysis: LH-P, DJA, JRG; Writing - original draft preparation: LH-P; Writing - review and editing: LH-P, MI, DJA, JRG, MRC.

## Supplementary information

There are ten supplementary figures, contained in the Supplementary Figures file, and eight supplementary tables, contained in the Supplementary Tables file.

## Data availability

Raw Xenium data is deposited on GEO. Images, processed data (including annotated, ready-to-use Seurat objects), and a Source Data file containing raw values for various plots across Figs. 1-4, are available on Zenodo (doi: 10.5281/zenodo.12625999). All data will be made publicly available as of the date of publication. Any additional data will be made available by the corresponding author upon reasonable request.

## Code availability

Code used for processing Xenium data and all downstream analysis, including figure generation, is available on GitHub: https://github.com/neurogenetics/Xenium_Line61_spatial.

