## Supplementary Figures for "Imaging spatial transcriptomics reveals molecular patterns of vulnerability to pathology in a transgenic α-synucleinopathy model"

Liam Horan-Portelance[1], Michiyo Iba[1], Dominic J. Acri[1], J. Raphael Gibbs[2], Mark R. Cookson[1*]

[1] Cell Biology and Gene Expression Section, Laboratory of Neurogenetics, National Institute on Aging, National Institutes of Health, Bethesda, MD 20892, USA.
[2] Computational Biology Group, Laboratory of Neurogenetics, National Institute on Aging, National Institutes of Health, Bethesda, MD 20892, USA.

* Corresponding author: Mark R. Cookson, PhD, Laboratory of Neurogenetics, NIA, NIH, Building 35, Room 1A116, MSC3707 35, Convent Drive, Bethesda, MD, 20892-3707, USA. Phone +1-301-451-3870.

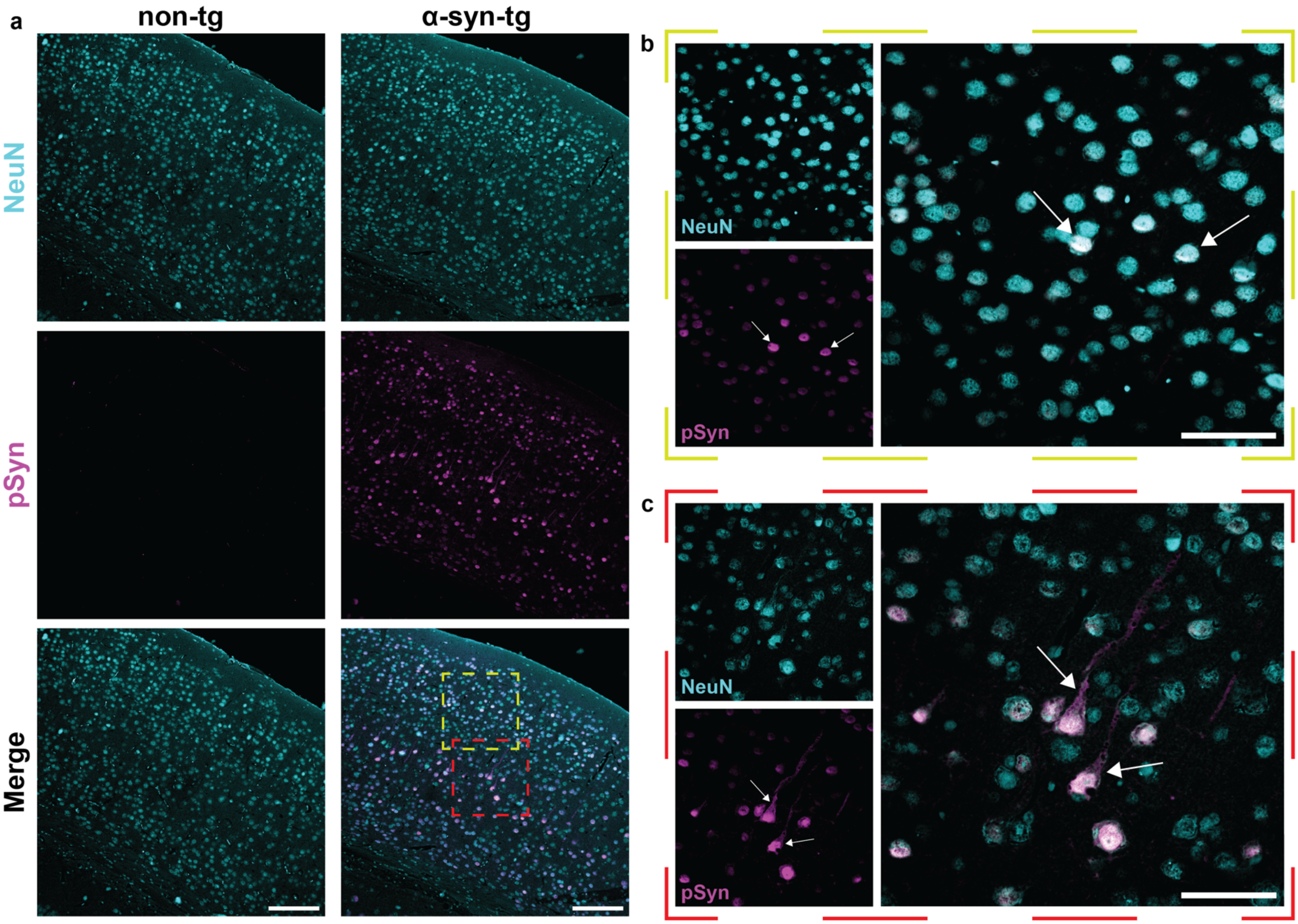


**Supplementary Fig. 1: Representative images of pSyn pathology in non-tg and α-syn-tg brains.** Following the Xenium run, sections were stained for NeuN (neuronal marker) and α-syn phosphorylated at Ser129 (pSyn). **a** Representative overview (10x magnification) images of non-tg (left) and α-syn-tg (right) cortices, showing staining of pan-neuronal marker NeuN (top, cyan), pSyn (middle, magenta), and merged. Images display absence of $pSyn^{+}$ cells in the non-tg cortex, and abundant neuronal pSyn staining in the α-syn-tg brain. Scale bar = 150 µm. **b** Representative high-magnification (40x) image of outer-layer (layer 2/3) neurons from an α-syn-tg cortex, showing neurons stained with NeuN (top left, cyan) and pSyn (bottom left, magenta), and merged (right). pSyn staining patterns highlight the nuclear localization of pathology in these cells (arrows). Scale bar = 50 µm. **c** Representative high-magnification (40x) image of deep-layer (layer 5) neurons from an α-syn-tg cortex, showing neurons stained with NeuN (top left, cyan) and pSyn (bottom left, magenta), and merged (right). pSyn staining patterns highlight the severe pathology in these cells extending out of the nucleus, to the cytoplasm and axons (arrows). Scale bar = 50 µm.

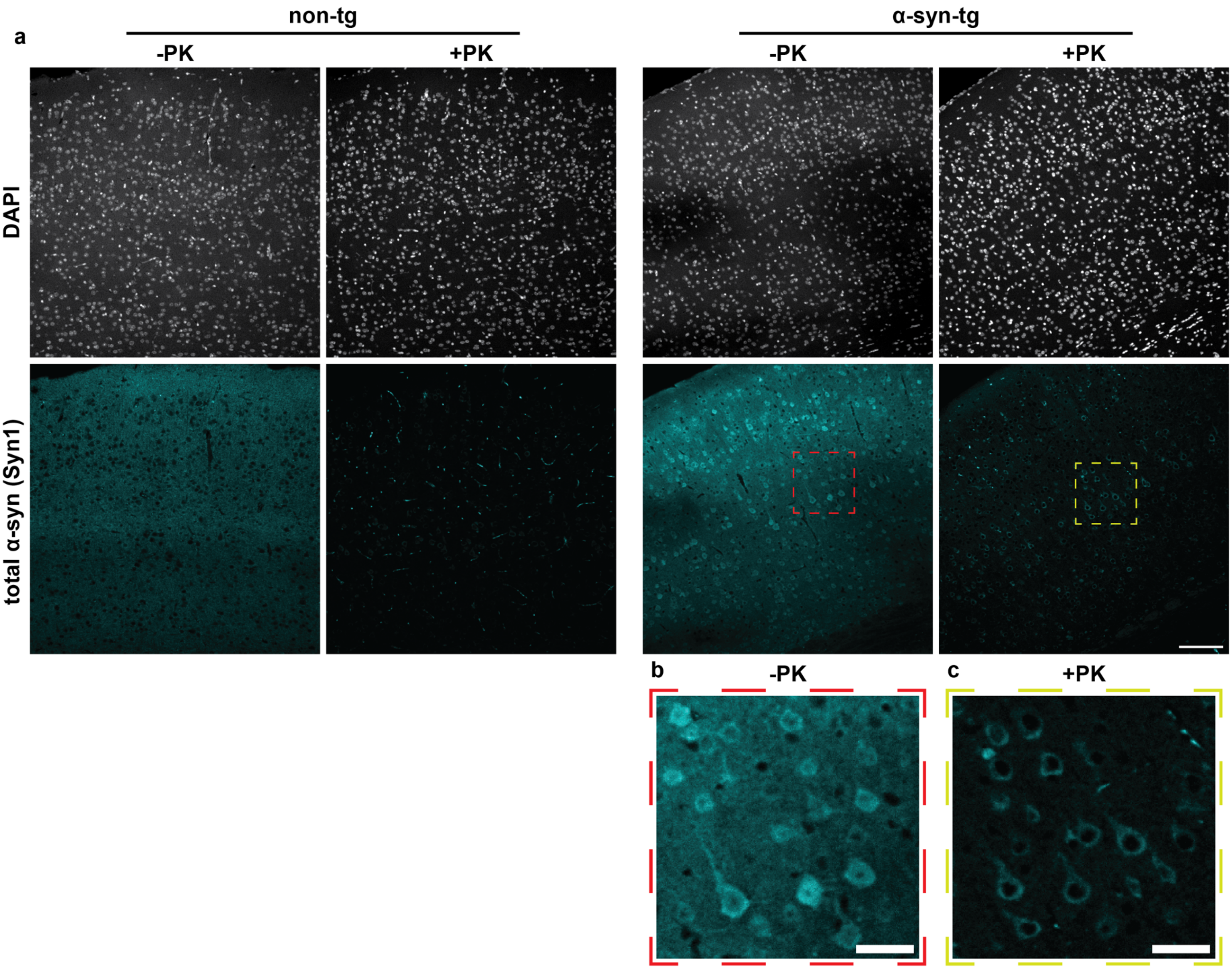


**Supplementary Fig. 2: Representative images of proteinase K-resistant α-synuclein pathology in non-tg and α-syn-tg brains. a** Representative immunofluorescence images of non-tg and α-syn-tg FFPE sections stained with an antibody against total α-syn. Sections were incubated in the presence of absence of proteinase K (PK) prior to staining. Images shown for non-tg and α-syn-tg (+/- PK) are from serial sections from the same animals. Scale bar = 100 µm. **b-c** Zoomed images of neurons in the cortex of an α-syn-tg animal incubated with and without PK, showing some PK-resistant α-syn remaining in the cell bodies of some neurons. Scale bar = 30 µm.

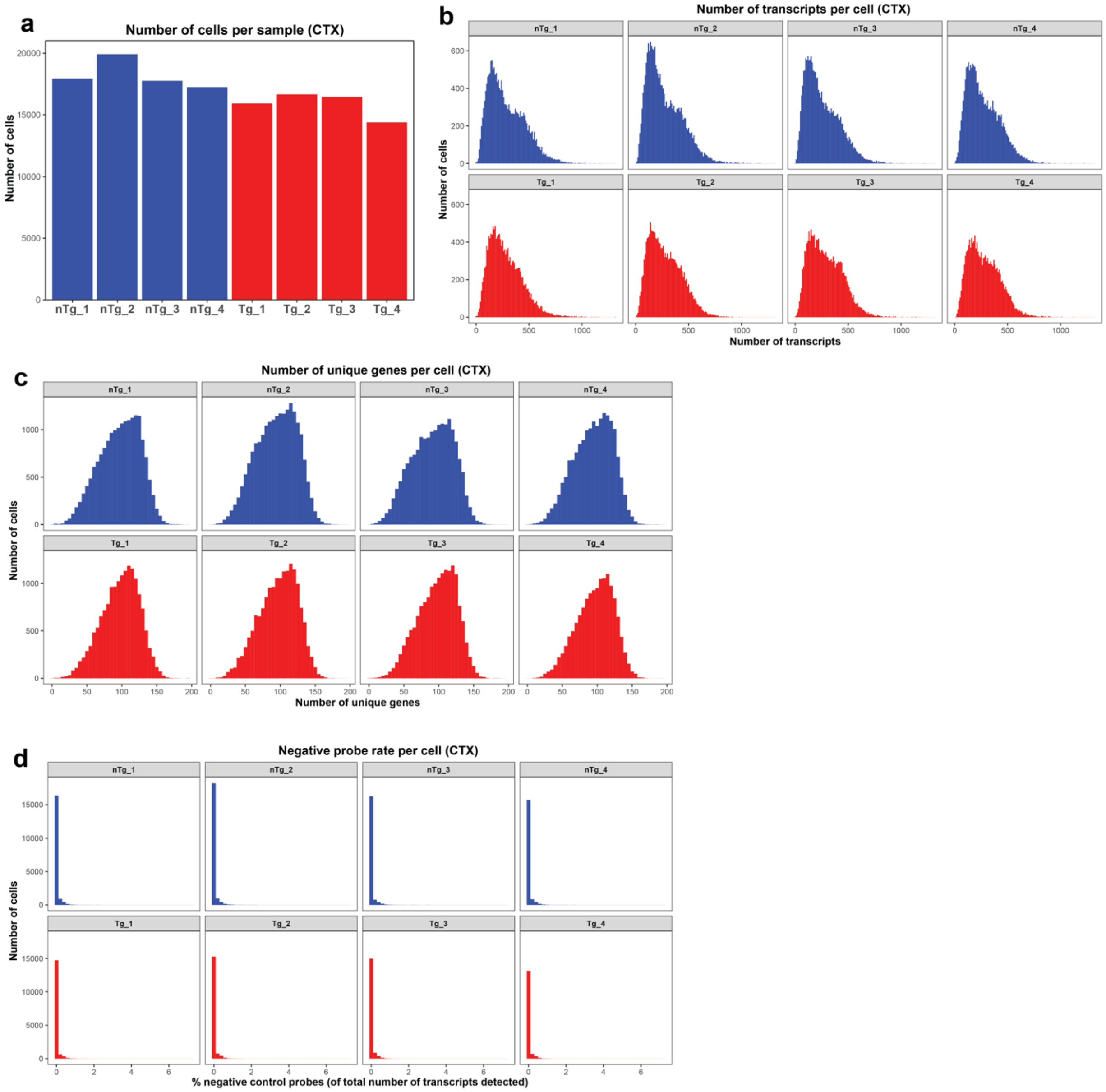


**Supplementary Fig. 3: Xenium quality control measures for cortical cells. a** Number of cells detected in each sample. **b** Number of transcripts detected per cell in each sample. **c** Number of unique genes detected per cell in each sample. **d** Negative probe rate for each cell in each sample, calculated as the number of detected negative control probes divided by the total number of detected probes in a given cell.

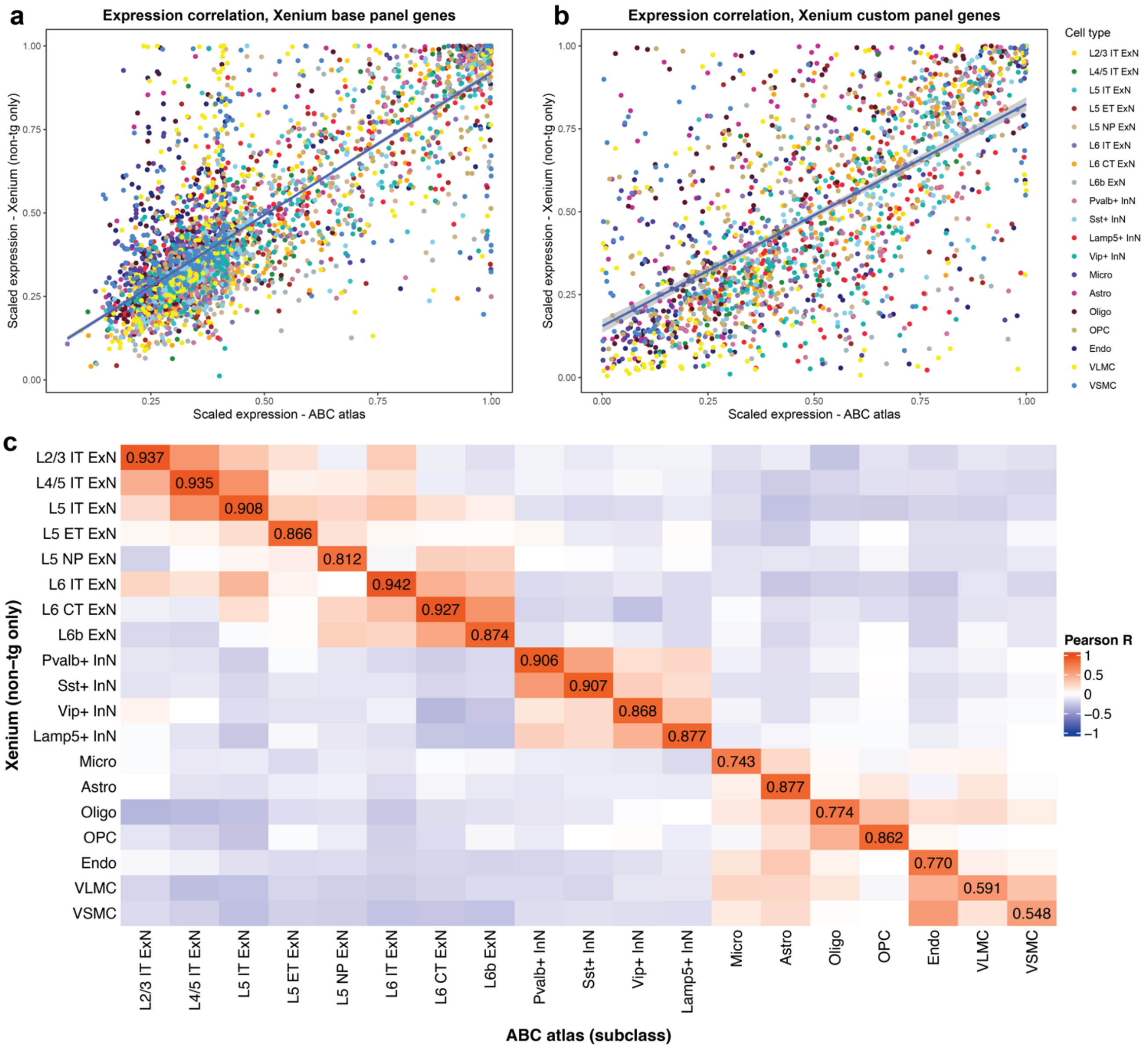


**Supplementary Fig. 4: Gene expression and cell type agreement between sequencing-based and imaging-based transcriptomics. a-b** Correlation of expression of the 248 genes in the Xenium base panel **(a)** and the 99 genes in our custom panel (excluding *hSNCA*) **(b)** between our Xenium data and the Allen Brain Cell atlas scRNA-seq data. Expression for each gene was scaled across cell types to allow for direct comparison, and each point represents scaled expression of a single gene in a single cell type (see Methods, "Comparison of Xenium to single-cell RNA sequencing data"). **c** Heatmap showing Pearson correlation of expression profiles between cell types, in Xenium and the ABC atlas, considering only the Xenium base panel genes. Scaled gene expression profiles (i.e., scaled expression of every gene in the base panel) for each cell type across both technologies were compared, and a Pearson coefficient was calculated between every cell type (see Methods, "Comparison of Xenium to single-cell RNA sequencing data"). Pearson R is displayed for each matching cell type pair.

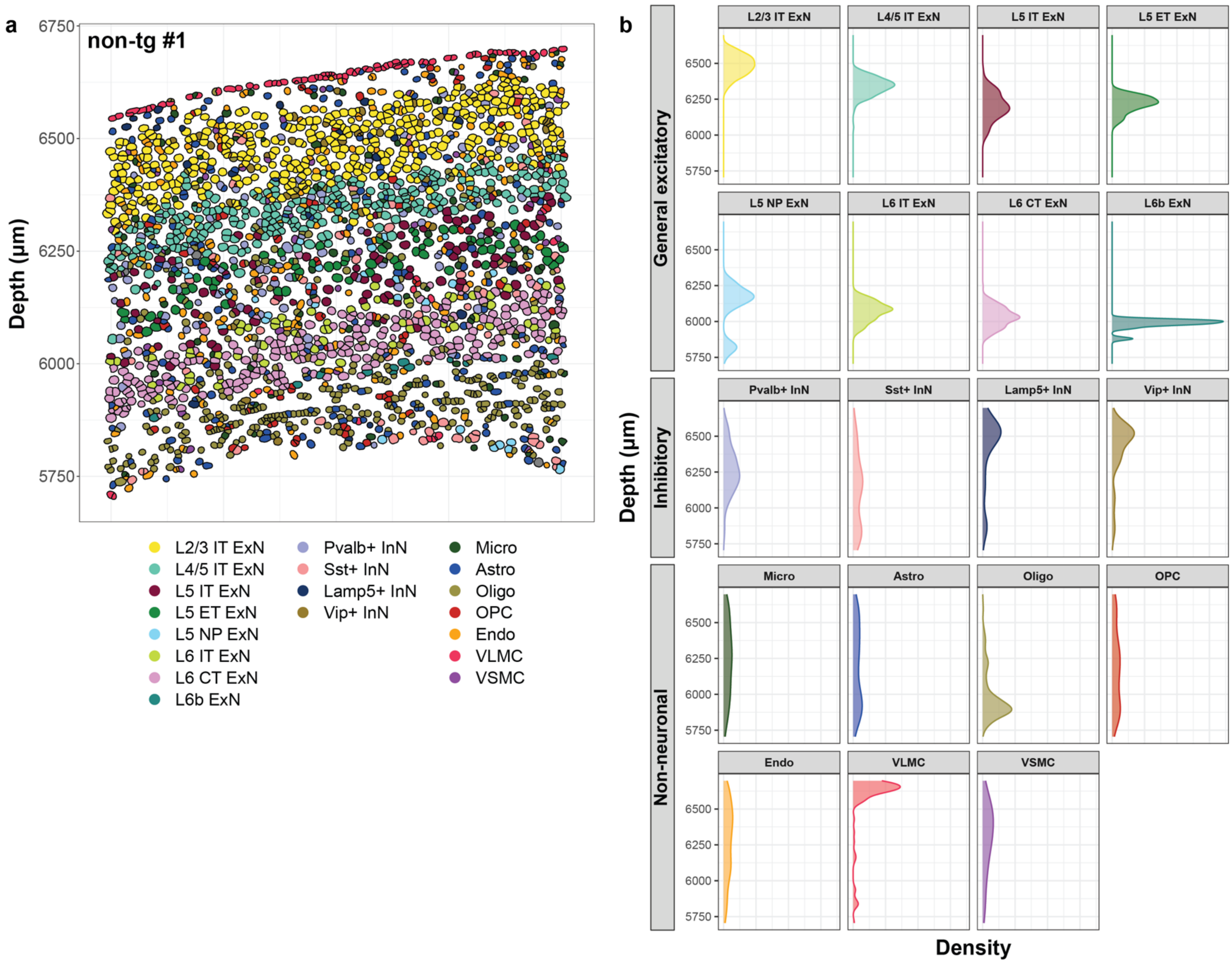


**Supplementary Fig. 5: Ground truth spatial information further validates Xenium cell type assignments in a non-tg brain. a** Representative image containing all 6 cortical layers and the corpus collosum (same as seen in Fig. 1b, from animal non-tg #1). Each dot represents a cell, colored by cell type. Depth is indicated to the left, represented as the x-coordinate (in µm). **b** Density plots showing distribution of each of the cell types as a function of depth through the cortex in the representative FOV in **(a)**.

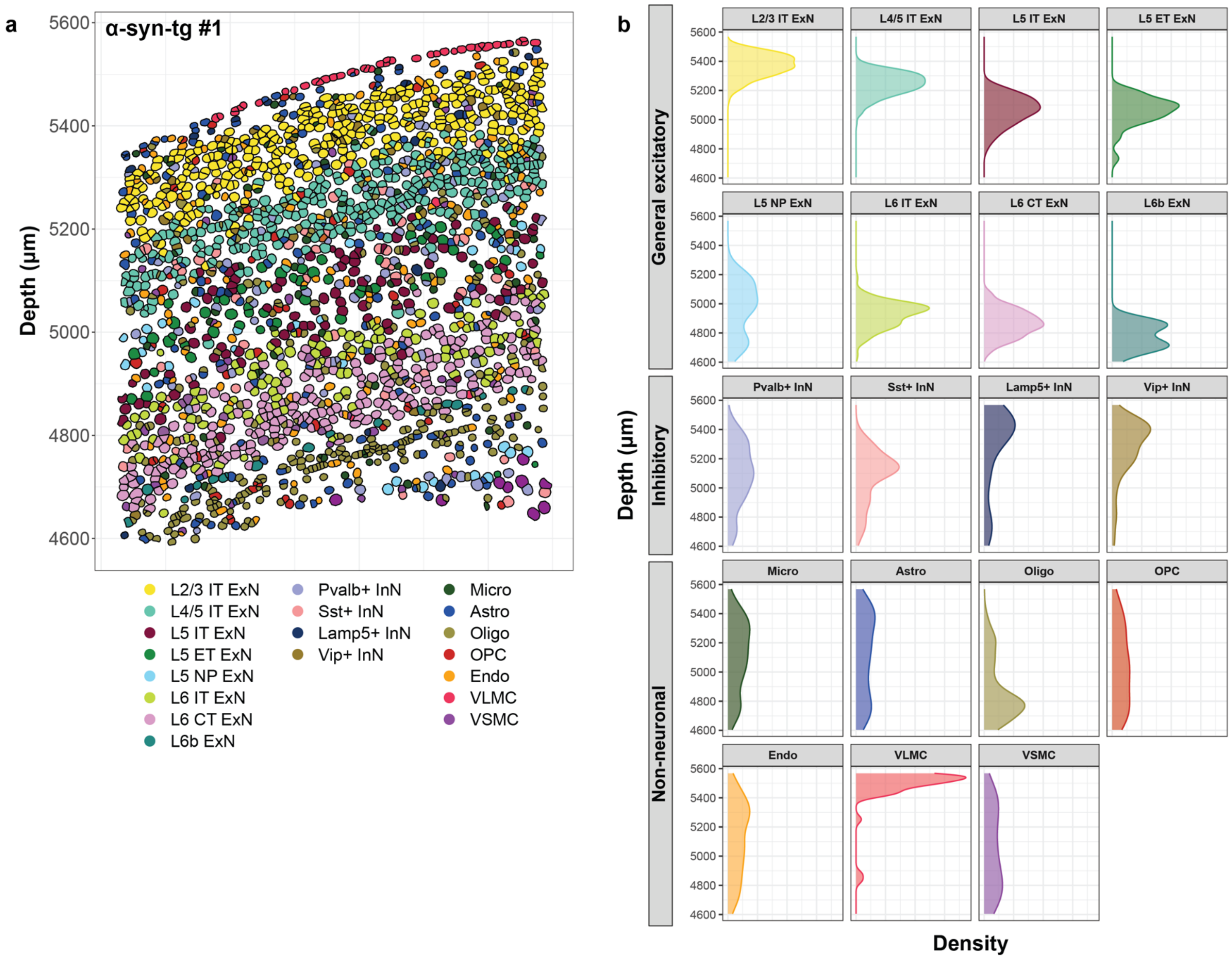


**Supplementary Fig. 6: Ground truth spatial information further validates Xenium cell type assignments in a an α-syn-tg brain. a** Representative image containing all 6 cortical layers and the corpus collosum (from animal α-syn-tg #1). Each dot represents a cell, colored by cell type. Depth is indicated to the left, represented as the x-coordinate (in µm). **b** Density plots showing distribution of each of the cell types as a function of depth through the cortex in the representative FOV in **(a)**.

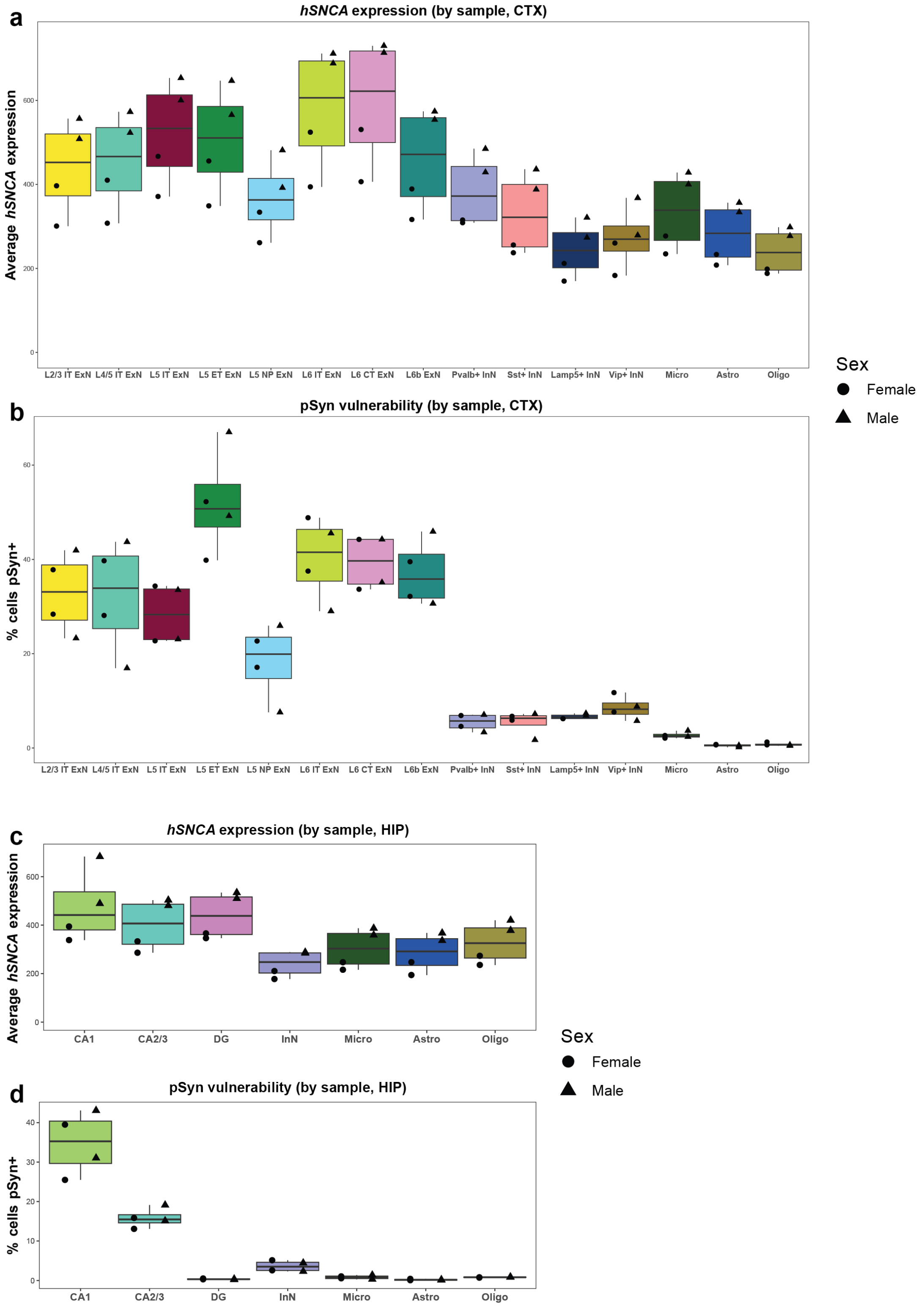


**Supplementary Fig. 7: α-Syn-tg mice show sex differences in *hSNCA* expression, but not pSyn pathology. a-d** *hSNCA* expression **(a, c)** and pSyn pathology frequencies **(b, d)** shown for the cortex **(a, b)** and hippocampus **(c, d)** in α-syn-tg mice. Each point represents one animal, with males shown as triangles and females shown as circles.

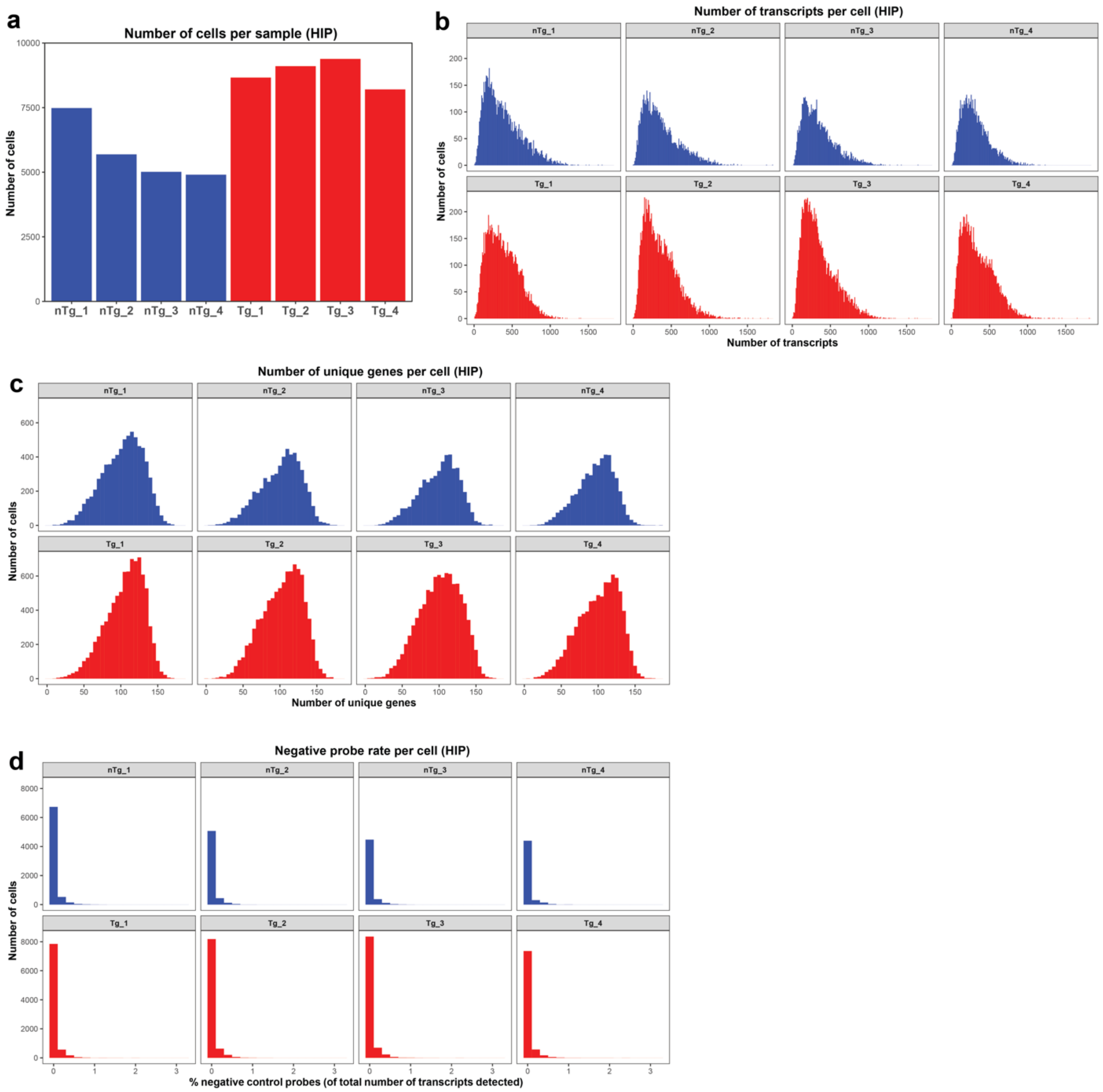


**Supplementary Fig. 8: Xenium quality control measures for hippocampal cells. a** Number of cells detected in each sample. **b** Number of transcripts detected per cell in each sample. **c** Number of unique genes detected per cell in each sample. **d** Negative probe rate for each cell in each sample, calculated as the number of detected negative control probes divided by the total number of detected probes in a given cell.

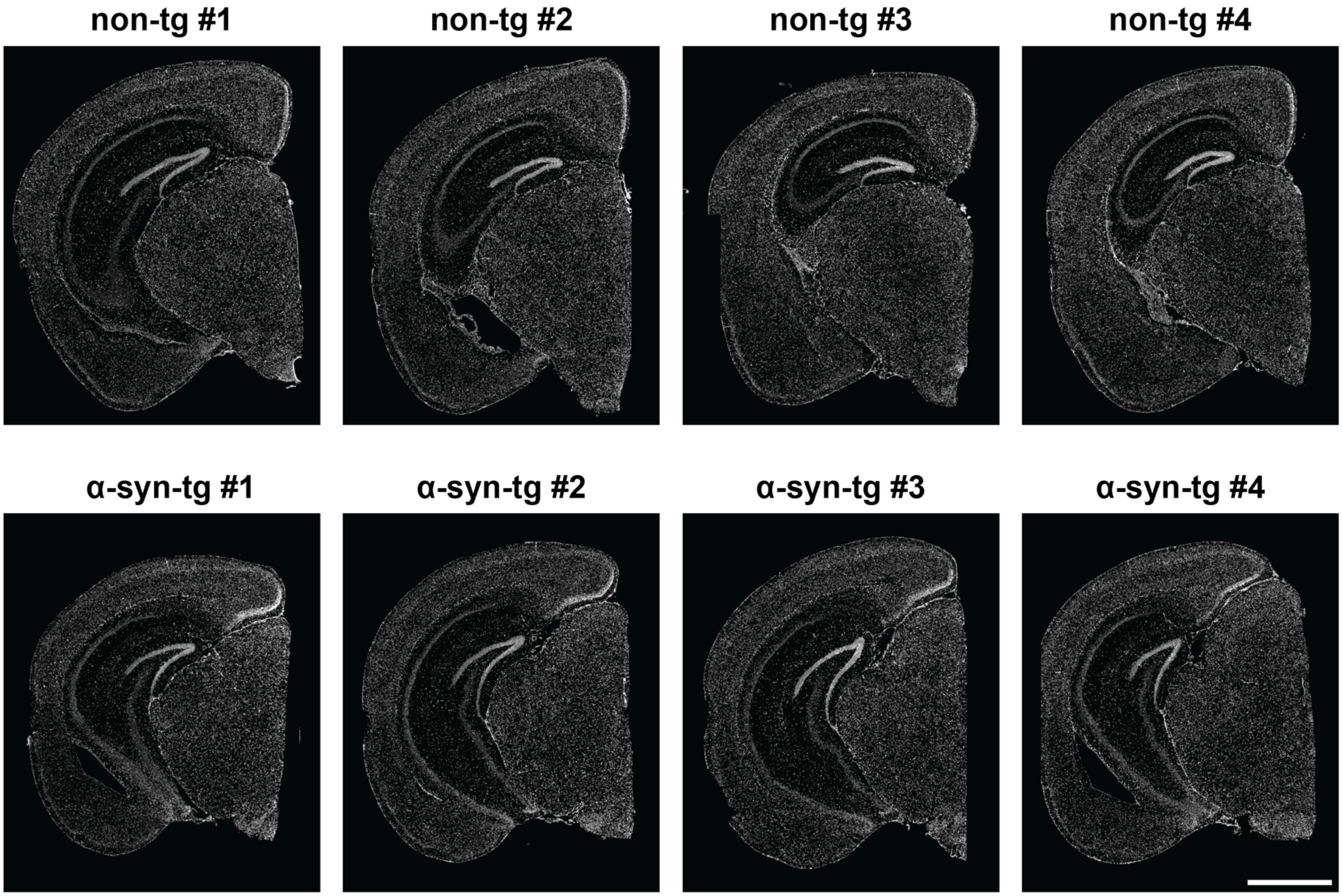


**Supplementary Fig. 9: Full images of all eight brains used for Xenium.** DAPI staining of all eight brains, images taken on the Xenium instrument. Scale bar = 1.5 mm.

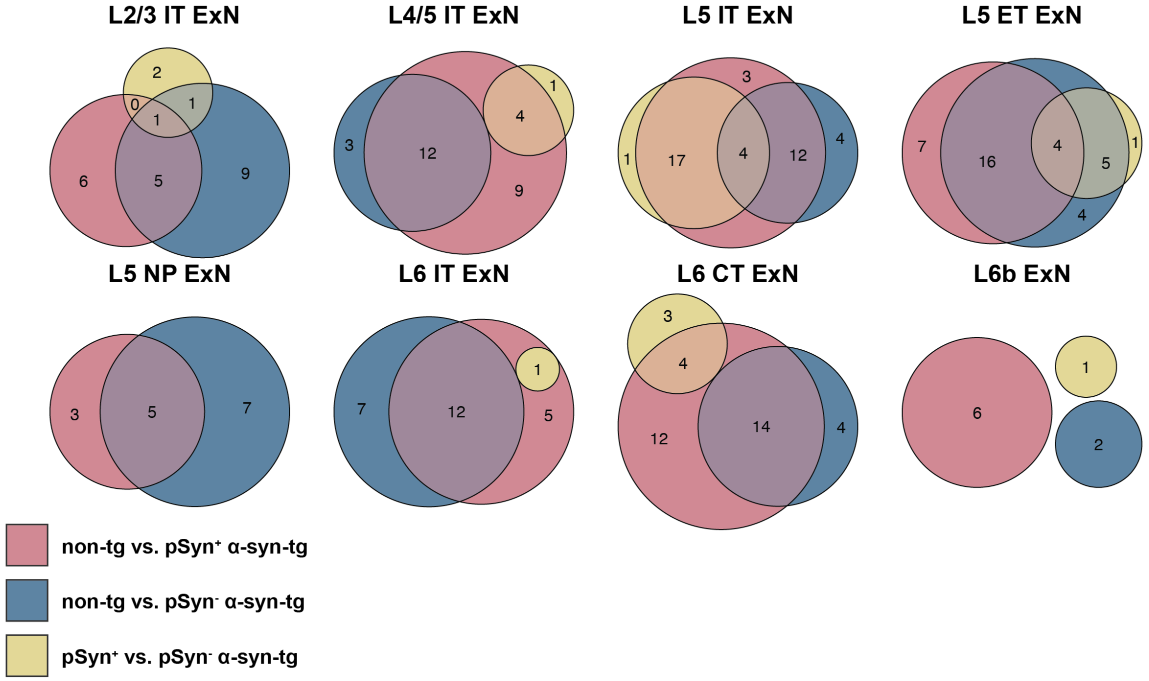


**Supplementary Fig. 10: DEG overlap across cell types between pseudo-bulk comparisons.** Euler plots for all general ExN subtypes showing the overlap of DEGs between the 3 differential expression comparisons (red = non-tg vs. pSyn$^{+}$ α-syn-tg; blue = non-tg vs. pSyn$^{-}$ α-syn-tg; yellow = pSyn$^{+}$ vs. pSyn$^{-}$ α-syn-tg). Size of the circles is scaled to the number of genes in each group, which is denoted by the numbers inside the circles.

**Captions for Supplementary Tables**

**Supplementary Table 1:** Information on the animals used in this study (animal ID, sex, age).

**Supplementary Table 2:** Genes included in the base and custom Xenium panels (gene name and Ensembl ID).

**Supplementary Table 3:** Counts of cell types in the cortex and hippocampus by animal.

**Supplementary Table 4:** Sequences for human *SNCA* and mouse *Snca* custom Xenium probes.

**Supplementary Table 5:** Linear regression results for four comparisons, % cells pSyn$^{+}$ vs.: non-tg GEx (all cell types), non-tg GEx (ExNs only), α-syn-tg GEx (all cell types), and α-syn-tg GEx (ExNs only).

**Supplementary Table 6:** All significant differentially expressed genes for pSyn$^{+}$/*hSNCA*-low vs. pSyn$^{-}$/*hSNCA*-high cells in the cortex (corresponds to main Fig. 5c).

**Supplementary Table 7:** All significant differentially expressed genes for pSyn$^{+}$ vs. pSyn$^{-}$ α-syn-tg cells in the hippocampus.

**Supplementary Table 8:** Full differential expression and GLM results, significant and non-significant (corresponds to main Fig. 6b-e).
